# Integrative multi-environmental genomic prediction in apple

**DOI:** 10.1101/2024.06.20.599822

**Authors:** Michaela Jung, Carles Quesada-Traver, Morgane Roth, Maria José Aranzana, Hélène Muranty, Marijn Rymenants, Walter Guerra, Elias Holzknecht, Nicole Pradas, Lidia Lozano, Frédérique Didelot, François Laurens, Steven Yates, Bruno Studer, Giovanni AL Broggini, Andrea Patocchi

## Abstract

Genomic prediction for multiple environments can aid the selection of genotypes suited to specific soil and climate conditions. Methodological advances allow effective integration of phenotypic, genomic (additive, non-additive), and large-scale environmental (enviromic) data into multi-environmental genomic prediction models. These models can also account for genotype-by-environment interaction, utilize alternative relationship matrices (kernels), or substitute statistical approaches with deep learning. However, the application of multi-environmental genomic prediction in apple remained limited, likely due to the challenge of building multi-environmental datasets and structurally complex models. Here, we applied efficient statistical and deep learning models for multi-environmental genomic prediction of eleven apple traits with contrasting genetic architectures by integrating genomic- and enviromic-based model components. Incorporating genotype-by-environment interaction effects into statistical models improved predictive ability by up to 0.08 for nine traits compared to the benchmark model. This outcome, based on Gaussian and Deep kernels, shows these alternatives can effectively substitute the standard G-BLUP. Including non-additive effects slightly improved predictive ability by up to 0.03 for two traits, but enviromic-based effects resulted in no improvement. The deep learning approach achieved the highest predictive ability for three traits with simpler genetic architectures, outperforming the benchmark by up to 0.10. Our results demonstrate that the tested statistical models capture genotype-by-environment interactions particularly well, and the deep learning models efficiently integrate data from diverse sources. This study will foster the adoption of multi-environmental genomic prediction to select apple cultivars adapted to diverse environmental conditions, providing an opportunity to address climate change impacts.

## INTRODUCTION

Since the introduction of genomic selection (Meuwissen et al., 2001), the genome-wide selection based on thousands of markers has resulted in increased genetic gain, and this approach is progressively becoming an integral component of modern crop breeding programs (Garcí a-Ruiz et al., 2016; Voss-Fels et al., 2019). To predict the genomic estimated breeding values for genomic selection, marker effects are frequently estimated using the well-established genomic best linear unbiased predictor (G-BLUP) approach (VanRaden, 2008). For genomic prediction across environments, increased predictive ability has been demonstrated by utilizing G-BLUP to incorporate the main marker effects and interaction effects of markers and environments (Jarquí n et al., 2014; Lopez-Cruz et al., 2015). The interaction between markers and environments provides a mathematical representation of the natural phenomenon of genotype-by-environment interaction, which results from the variability in the genotype performance ranking across different environmental conditions. Despite numerous reports of successful phenotypic performance prediction using molecular markers in perennial crops such as apple (Kostick et al., 2023; Kumar et al., 2012; Migicovsky et al., 2016; Muranty et al., 2015), genotype-by-environment interaction has been often overlooked in genomic prediction of apple traits.

The most comprehensive study conducted thus far to investigate the influence of genotype-by-environment interaction on genomic predictive ability in apple, conducted by Jung et al. (2022), was achieved by the establishment of the apple reference population, known as the apple REFPOP (Jung et al., 2020). Across the numerous phenotypic traits assessed in the apple REFPOP, genotype-by-environment interaction explained up to 24% of the phenotypic variance, and the incorporation of genotype-by-environment interaction into G-BLUP resulted in a predictive ability increase of up to 0.07 (Jung et al., 2022). The challenge of building multi-environmental datasets, coupled with the computational costs tied to the structural complexity of genomic prediction models accommodating genotype-by-environment interaction, has likely limited the use of such models in practice.

Recent software advances that reduce computational time could enable broader adoption of multi-environmental genomic prediction models in plant breeding (Costa-Neto, Fritsche-Neto, et al., 2021; Granato et al., 2018). Empirical comparisons between the well-established R package ‘BGLR’ (Pe rez & de los Campos, 2014) and the newer R package ‘BGGE’ (Granato et al., 2018), both of which apply the same model structures based on G-BLUP, revealed comparable predictive abilities, but ‘BGGE’ was up to five times faster (Granato et al., 2018). In addition to G-BLUP, covariance matrices, alternatively referred to as relationship matrices or kernels, can be estimated using approaches that capture nonlinearity in the relationships between phenotype and genotype. The nonlinear Gaussian kernel and the Deep kernel (also known as the arc-cosine kernel) have demonstrated superior performance compared to G-BLUP, showing reduced computational time and increased predictive ability in maize and wheat datasets (Costa-Neto, Fritsche-Neto, et al., 2021; Cuevas et al., 2019).

In addition to the commonly used genomic effects of molecular markers, the advancements in software have introduced straightforward options for incorporating additional sources of variation into genomic prediction models (Costa-Neto, Fritsche-Neto, et al., 2021; Costa-Neto, Galli, et al., 2021). Marker values can be split into additive values and dominance deviations that allow for orthogonal partition of variances, which implies that the proportions of additive genomic effects remain constant even when dominance effects are incorporated into the genomic prediction model (A lvarez-Castro & Carlborg, 2007). The incorporation of additive and dominance effects into genomic prediction models is typically done by the use of covariance matrices, as proposed by Vitezica et al. (2013, 2017). A combination of these effects with a fixed effect of inbreeding has demonstrated improved genomic predictive ability in both maize and sugarcane (Roth et al., 2022; Yadav et al., 2021). However, in a perennial fruit crop such as blueberry, the inclusion of dominance effects did not affect predictive ability (Amadeu et al., 2020). Additionally, incorporating non-genetic effects derived from large-scale assessment of environmental attributes (i.e, envirotyping, resulting in environmental covariates also called enviromic markers (Cooper et al., 2014; Resende et al., 2021)) into genomic prediction models can improve the estimation of similarities between environments and genotype-by-environment interaction. This enhancement not only leads to increased predictive ability, but also offers a more comprehensive understanding of the complex interplay between genetic and environmental factors (Costa-Neto, Fritsche-Neto, et al., 2021; Jarquí n et al., 2014). The enviromic-based effects, as well as the marker-based effects expressed as standard genomic, orthogonal additive and dominance effects, can all be studied as extensions of G-BLUP using conventional statistical genomic prediction model frameworks, which simplifies their integration into the modeling process.

Deep learning approaches have emerged as an alternative to conventional statistical genomic prediction models. The literature review of Montesinos-Lo pez et al. (2021) on the application of deep learning for genomic selection showed no distinct superiority of deep learning approaches in terms of predictive ability compared to conventional genomic prediction models, unless very large datasets were used. However, deep learning models allow for effective integration of data from diverse sources, but they can also become impractical for datasets containing many variables, leading to computational complexity and overfitting. In plant breeding, datasets comprising thousands of markers are compiled, and dimensional reduction may help simplify marker information for deep learning (Kick et al., 2023). In the study by Jurado-Ruiz et al. (2023), the use of a small subset of associated markers was critical for accurate predictions of apple shape when deploying neural networks. The potential application of deep learning for multi-environmental genomic prediction of diverse quantitative apple traits has yet to be examined.

This study aims to conduct a comprehensive comparison between conventional statistical models that integrate genomic- and enviromic-based effects and a deep learning approach for multi-environmental genomic prediction of apple traits. The subjects of prediction were eleven quantitative traits related to phenology, productivity, and fruit quality, which were measured from the apple REFPOP during five years at up to five locations, i.e., up to 25 environments (defined as combinations of location and year). The increased extent of the apple REFPOP dataset across environments allows an evaluation of different modeling techniques to harness the full potential of these data for accurate prediction of phenotypic traits. The main objectives of the study were: (i) to compare the relative contribution of different model components, i.e., random effects and feature streams, between the statistical and deep learning genomic prediction models, (ii) to assess and compare predictive abilities of these models, (iii) to evaluate and compare the computational time required for model training and prediction. By addressing these three crucial factors, this research aims to provide insights into the strengths and limitations of statistical models and deep learning to identify the best modelling solutions for the selection of apple cultivars adapted to diverse environmental conditions.

## RESULTS

### Dataset composition

From the eleven phenotypic traits assessed in the apple REFPOP over five years and at a maximum of five locations, two environment-trait combinations were excluded due to very low values of the environment-specific clonal mean heritability (*H*^2^ < 0.1). The excluded combinations included phenotypic measurements for floral emergence in Spain in 2020 (*H*^2^ = 0.036) and flowering intensity in France in 2021 (*H*^2^ = 0.002). Consequently, phenotypic estimates were generated from a minimum of eight environments for titratable acidity, soluble solids content, and fruit firmness, while harvest date, total fruit weight, number of fruits, and single fruit weight were evaluated across the maximum number of environments, totaling 25 (Table S1). Various shapes of distributions and consistent patterns of Pearson’s correlations were observed for the adjusted means of phenotypic traits over years and locations (Figure 1A, Figure S1, Figure S2).

**Figure 1:**
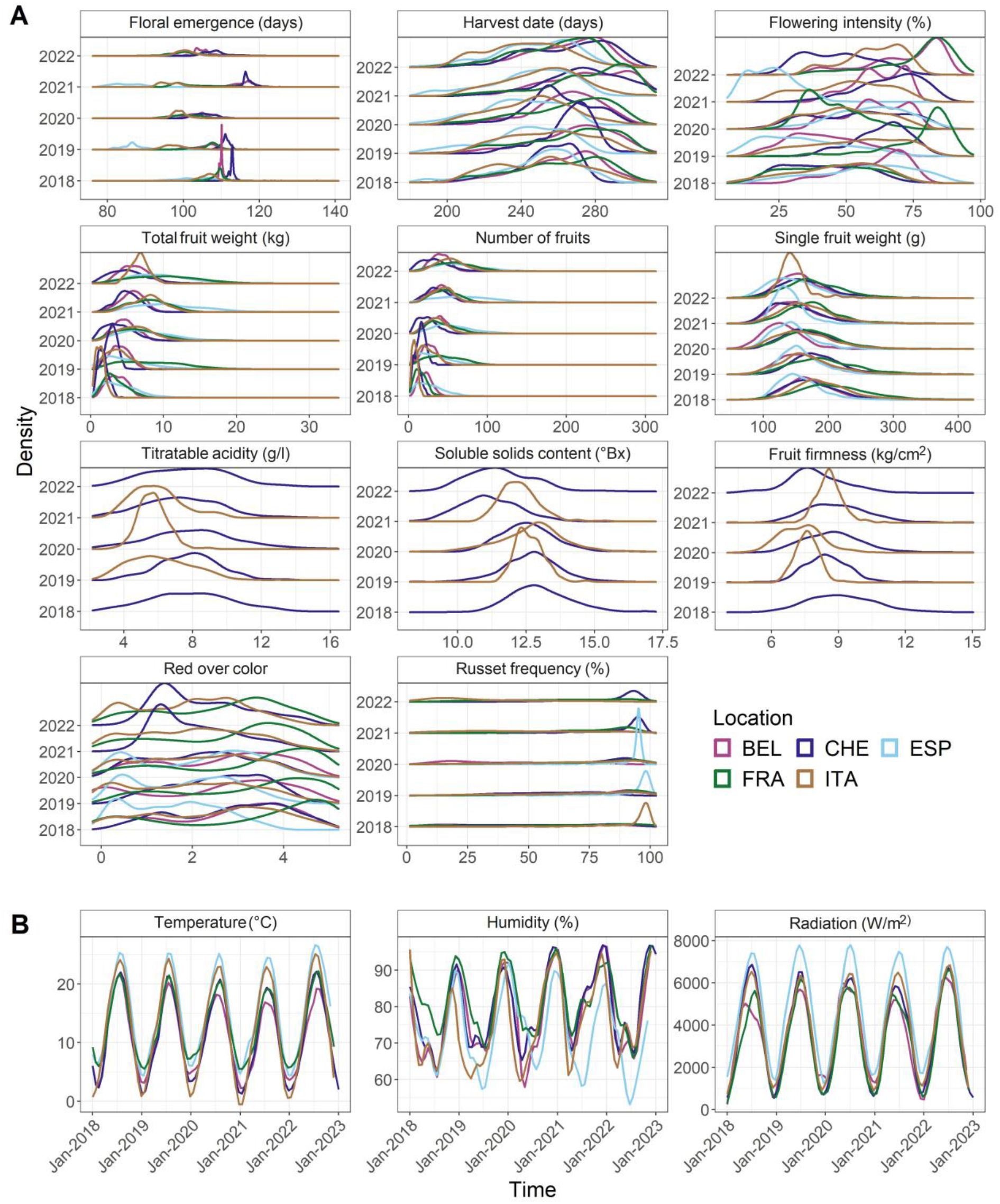
Phenotypic and weather data distributions. **A** Density estimates for the adjusted means of eleven phenotypic traits from five locations and five years of measurement. The locations correspond to Belgium (BEL), Switzerland (CHE), Spain (ESP), France (FRA) and Italy (ITA). **B** Local regression curves spanning five years estimated from daily temperature means, daily humidity means and daily radiation sums. Colors correspond to legend in A.

For the weather variables, moderate differences were observed in daily temperature means, daily humidity means, and daily radiation sums between years and locations (Figure 1B). Consequently, these data were summarized based on phenology, meaning the data was split into two periods: the first 80 days until 90% of the genotypes flowered, and the following days until 90% of the genotypes were harvested (Figure S3). After preprocessing the soil variables, the final enviromic dataset included 28 environmental covariates for weather and soil.

### Relationship matrices

Implementation of the G-BLUP approach resulted in the standard genomic relationship matrix *K*_*G*_ (based on standard allele coding with allele dosage values of 0, 1, and 2), the additive genomic relationship matrix *K*_*A*_, and the dominance genomic relationship matrix *K*_*D*_ (Figure 2A, B and C). The heatmaps of these matrices depicted a strong similarity between *K*_*G*_ and *K*_*A*_ (Figure 2A and B). The lower-left quadrant of matrices *K*_*G*_ and *K*_*A*_ comprised the apple REFPOP accessions, revealing only subtle differences between these genotypes. The upper-right quadrant of matrices *K*_*G*_ and *K*_*A*_ visualized the apple REFPOP progenies grouped according to their biparental origin. The progeny groups were evident in the matrix *K*_*D*_, but no further strong relationships between genotypes were visually observed. *K*_*A*_ and *K*_*D*_ showed the mean of their matrix values close to zero and the mean of the diagonal of 1. Gaussian kernel and Deep kernel, used as alternative approaches to G-BLUP, resulted in matrices *K*_*GGK*_ and *K*_*GDK*_ (Figure 2D and E) that were visually similar to the *K*_*G*_ and *K*_*A*_ matrices implemented using G-BLUP (Figure 2A and B), although some differences were observed particularly for the Gaussian kernel approach (Figure 2D). Application of the G-BLUP to the enviromic dataset of 28 environmental covariates resulted in the enviromic relationship matrix *K*_*W*_ (Figure 3). Hierarchical clustering of the matrix *K*_*W*_ showed five clusters of environments, each cluster referring to one of the orchard locations.

**Figure 2:**
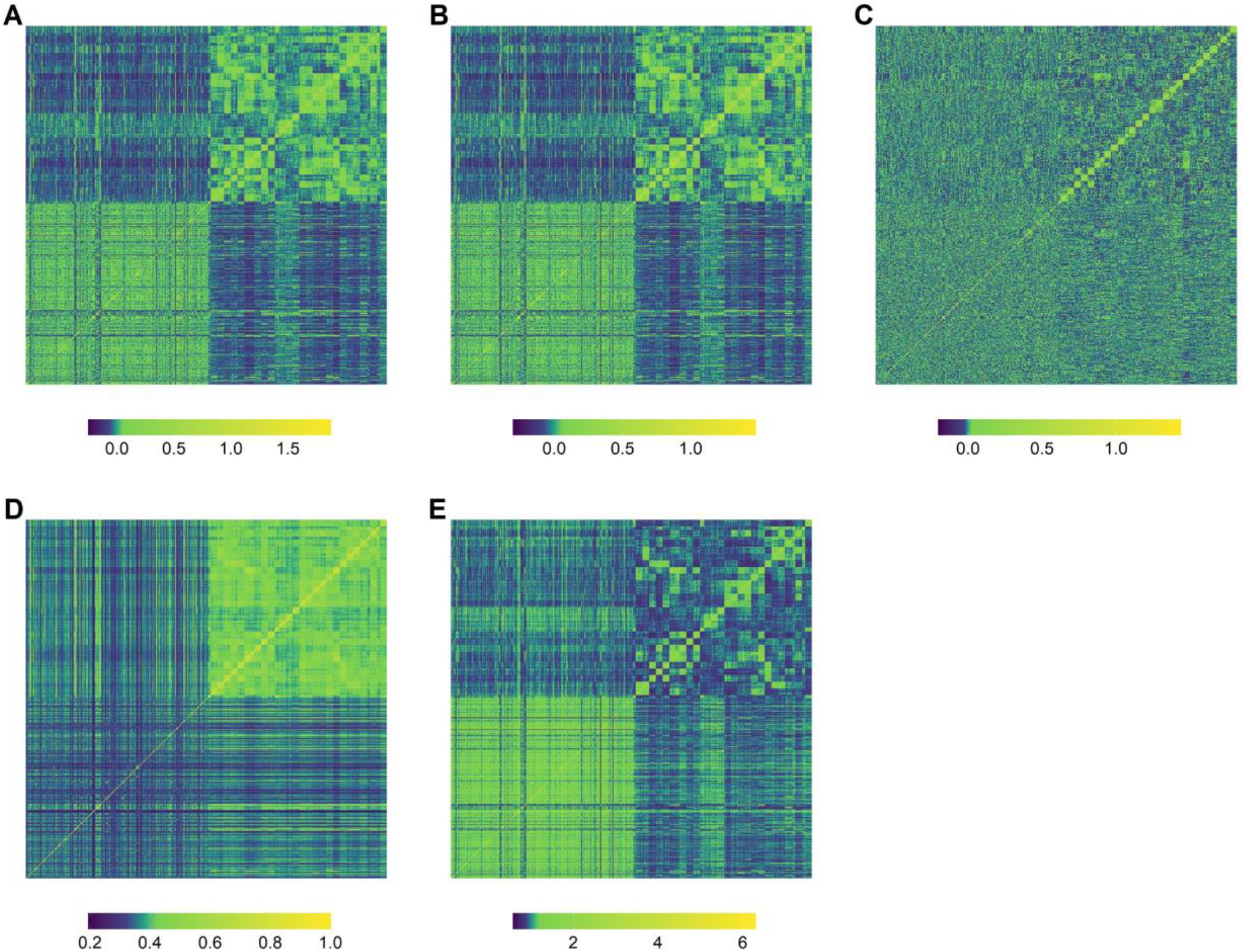
Heatmaps of the genomic relationship matrices. **A** Standard genomic relationship matrix *K*_*G*_ based on a marker matrix using the standard coding for bi-allelic SNPs (allele dosage values of 0, 1, and 2). **B** Additive genomic relationship matrix *K*_*A*_ based on marker matrix using the additive coefficients. **C** Dominance genomic relationship matrix *K*_*D*_ based on marker matrix using the dominance coefficients. The matrices in A–C were constructed using the G-BLUP approach. **D** Standard genomic relationship matrix *K*_*G_GK_*_ constructed deploying the Gaussian kernel (GK). **E** Standard genomic relationship matrix *K*_*G_DK_*_ based on the Deep kernel (DK). The lower-left and upper-right quadrants show the apple REFPOP accessions and progenies, respectively.

**Figure 3:**
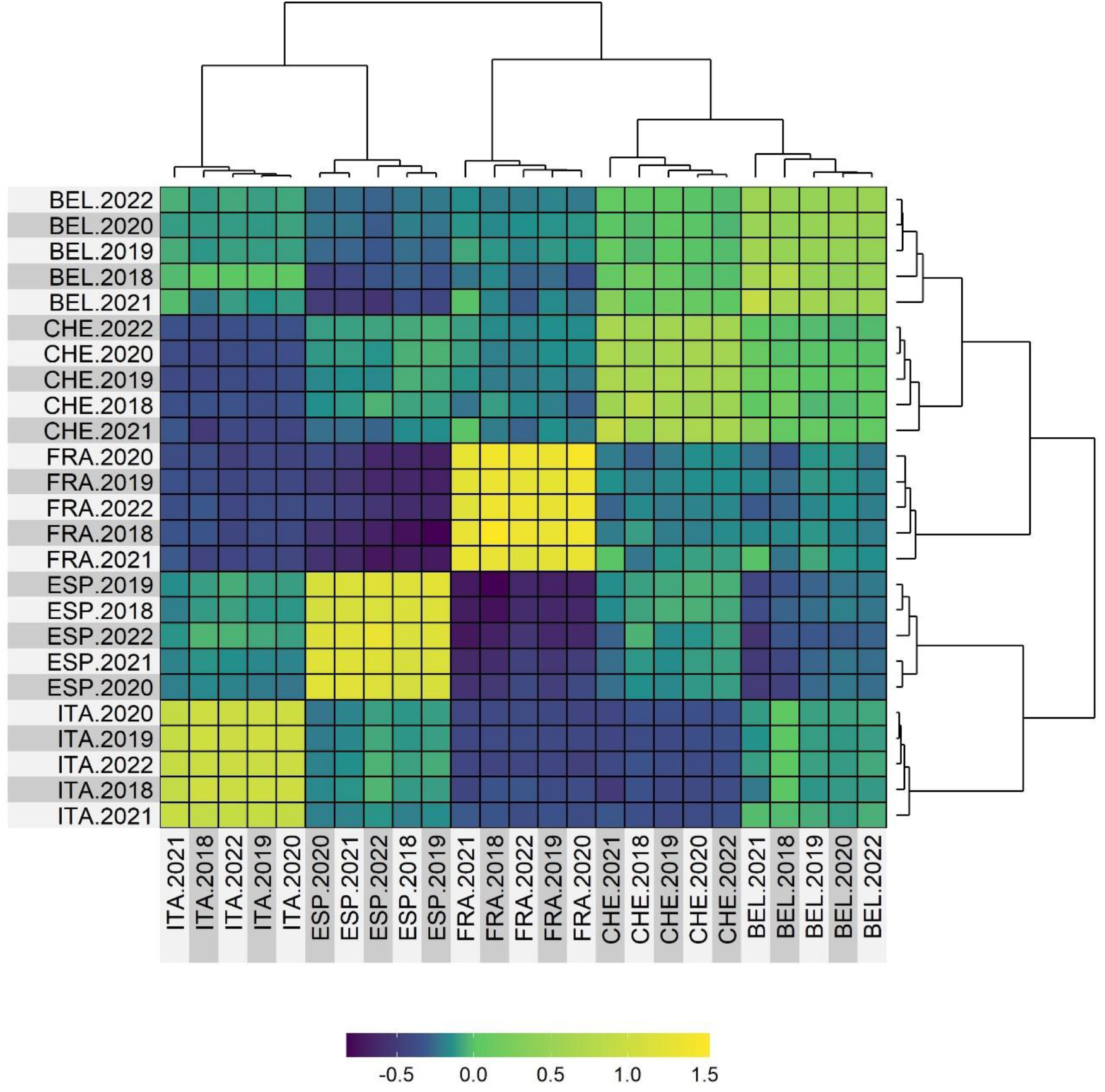
The enviromic relationship matrix *K*_*W*_ constructed from the environmental covariates for weather and soil using G-BLUP. Environments (combinations of location and year) were grouped applying hierarchical clustering.

### Contribution of the model components

Decomposition of the phenotypic variance using linear mixed models by incorporating random effects for the vector of genotypes (i.e., genotypic effects) and genotype-by-environment interaction, revealed that the proportion of phenotypic variance explained by the genotypic effects ranged from 9% for flowering intensity to 78% for harvest date (Figure 4A, Table S2). In contrast, the largest proportion of phenotypic variance explained by genotype-by-environment interaction was observed for flowering intensity (29%). The lowest proportion of genotype-by-environment interaction variance (9%) was found for harvest date. The total variance explained by both genotypic and genotype-by-environment interaction effects reached 64% on average across traits (Table S3).

**Figure 4:**
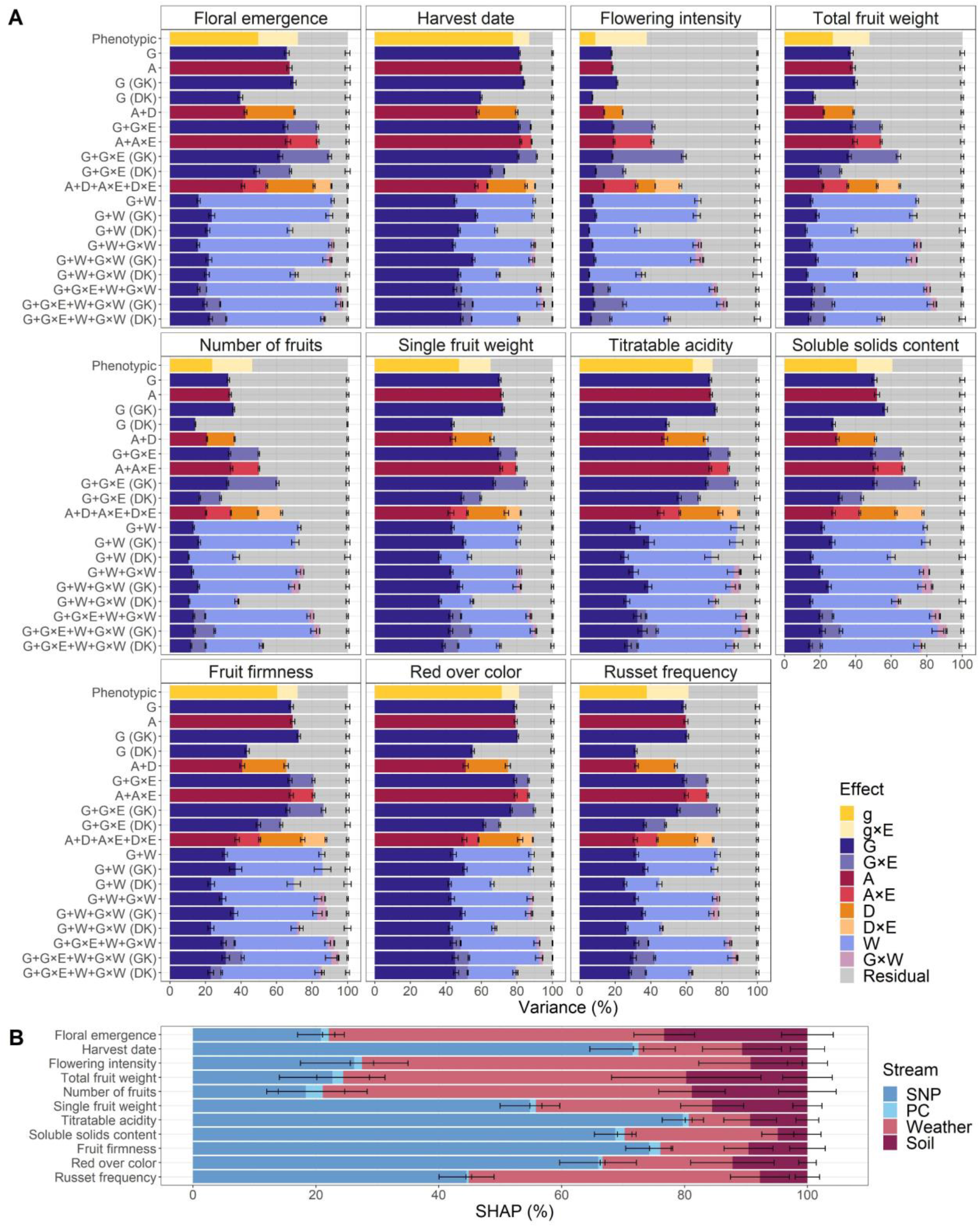
Relative contribution of different model components estimated for eleven traits. **A** Average proportions of phenotypic variance related to genotypic (g), genomic (G), additive (A) and dominance (D) effects, their interactions (×) with the vector of environments (E), the enviromic effects (W), the interaction effects G×W as well as the residual effect extracted from the statistical genomic prediction model fits. The relationship matrices for the different effects in the statistical genomic prediction models were constructed using the G-BLUP approach or, where indicated, the Gaussian kernel (GK) or deep kernel (DK). The statistical genomic prediction models were compared with a model based on phenotypic data (Phenotypic). Error bars correspond to standard deviation around the mean. **B** Relative contribution of the single nucleotide polymorphism (SNP), principal component (PC), weather, and soil feature streams estimated using Shapley additive explanations (SHAP) for the deep learning genomic prediction model. Error bars correspond to standard deviation around the mean.

For the statistical genomic prediction based on G-BLUP, linear mixed model structures resulted from the application of the relationship matrices *K*_*G*_, *K*_*A*_, and *K*_*D*_, representing genomic (G), additive (A), and dominance (D) effects, respectively. Various proportions of phenotypic variance related to these random effects and their interactions (×) were extracted from the model fits (Figure 4A, Table S2). Due to its model structure, the simplest genomic prediction model (used as a benchmark) was labeled as G, and its random genomic effects accounted for an average of 58% of the variance across traits (Table S3). Across all traits, model A explained ∼1% more variance compared to model G (Table S3). For model A+D, the average total proportion of variance explained by the model components A and D across traits was slightly lower than that of model G (difference of 0.3%, Table S3). Both models G+G×E and A+A×E, including interactions with the environment, explained, on average across traits, a proportion of variance 14% greater than that explained by model G (Table S3). Specifically, the effect G or A accounted for variance ranging from 18% for flowering intensity (G and A) to 81% (G) and 82% (A) for harvest date, and G×E and A×E explained variance ranging from 6% for harvest date (G×E and A×E) to 23% (G×E) and 22% (A×E) for flowering intensity (Table S2). The model A+D+A×E+D×E, on average across traits, explained a proportion of variance 21% and 7% greater than that explained by models G and G+G×E, respectively (Table S3).

The enviromic effects (W) and the interaction effects G×W were implemented applying the relationship matrix *K*_*W*_ based on G-BLUP in the model structures G+W, G+W+G×W, and G+G×E+W+G×W, and these models explained, on average across traits, 24%, 25%, and 30% more variance than model G, respectively (Figure 4A, Table S3). For the most complex model G+G×E+W+G×W, the proportions of variance explained by the interaction effects G×E and G×W were modest, ranging from 4% to 9% for G×E and 2% to 4% for G×W (Table S2).

When comparing models based on G-BLUP with their counterparts implementing Gaussian kernel using the relationship matrix *K*_*GGK*_ (model structures labeled with GK), the models G (GK), G+G×E (GK), and G+G×E+W+G×W (GK) demonstrated an average increase in explained variance of 3%, 7%, and 3% across traits, respectively (Figure 4A, Table S3). However, the models G+W (GK) and G+W+G×W (GK) resulted in an average decrease in explained variance of up to 1% (Table S3). On average over traits, the model structures based on Deep kernel implementing the relationship matrix *K*_*GDK*_ (model structures labeled with DK) exhibited a strong decrease in the proportion of variance explained by the genomic- and enviromic-based random effects when compared to their counterparts utilizing G-BLUP, namely -22% for G (DK), -19% for G+G×E (DK), -26% for G+W (DK), -25% for G+W+G×W (DK), and -17% for G+G×E+W+G×W (Table S3).

The applied deep learning genomic prediction model integrated marker and enviromic data through four feature streams, namely single nucleotide polymorphism (SNP), principal component (PC), weather, and soil streams, and the estimation of Shapley additive explanations (SHAP) revealed the relative mean importance of these feature streams (Figure 4B, Table S4). Across all traits, the relative SHAP contributions were 50% for the SNP stream, 1% for the PC stream, 36% for the weather stream, and 13% for the soil stream. The relative SHAP contribution for the SNP stream ranged from 18 to 26% for floral emergence and the productivity traits (flowering intensity, total fruit weight and number of fruits) to 80% for titratable acidity. For the PC stream, the relative SHAP contribution ranged between 0% for russet frequency and 3% for number of fruits. The lowest weather stream contribution of 10% was found for titratable acidity, while the largest contribution of the weather stream of 55 to 63% was found for floral emergence and the productivity traits (flowering intensity, total fruit weight and number of fruits). The relative SHAP contribution for the soil stream ranged between 5% for soluble solids content and 19 to 23% for floral emergence and two productivity traits (total fruit weight and number of fruits). An abundance of SNPs displaying high absolute mean SHAP were found for harvest date, titratable acidity and red over color (Figure S4, Figure S5). For harvest date, three SNPs with the highest absolute mean SHAP of 0.002 were located on chromosome 3 at 29.2 Mb (AX-115250472), 30.7 Mb (AX-115366114), and 30.8 Mb (AX-115233388). The three SNPs with the highest absolute mean SHAP of 0.003 for titratable acidity were found on chromosome 8 at 10.7 Mb (AX-115276534), 10.8 Mb (AX-115254093), and 11.8 Mb (AX-115519462). For red over color, the three SNPs with the highest absolute mean SHAP of 0.005 were located on chromosome 9 at 33.8 Mb (AX-105213720, AX-115558498), and 35.6 Mb (AX-115370846).

### Predictive ability

Assessment of genomic prediction model performance using five-fold cross-validation resulted in strong differences in predictive ability among the examined traits (Figure 5, Table S5). For the model G, the lowest predictive ability was observed for flowering intensity (0.25), russet frequency (0.32) and soluble solids content (0.37). Predictive abilities ranging from 0.41 to 0.51 were found for floral emergence, total fruit weight, number of fruits, single fruit weight, titratable acidity, and fruit firmness. The highest predictive ability of 0.69 was observed for harvest date, followed by the predictive ability of 0.63 for red over color. In comparison to the benchmark model G, model A demonstrated a significantly higher predictive ability of 0.64 for red over color. After a fixed effect of inbreeding was incorporated into model A, the resulting model A (inb) showed a significantly higher predictive ability than model G for the trait single fruit weight with an increase in predictive ability of 0.01 (Table S5). Model A+D (as well as model A+D (inb)) significantly outperformed model G for harvest date and red over color with a difference in predictive ability of 0.01 and 0.03, respectively. The incorporation of the G×E effect in model G+G×E yielded significantly higher predictive abilities compared to model G across multiple traits, including floral emergence, number of fruits, single fruit weight, soluble solids content, and russet frequency, with an increase in predictive ability ranging from 0.01 to 0.02. For flowering intensity, the G+G×E model resulted in a significant increase in predictive ability of 0.07. Model A+A×E significantly outperformed model G for floral emergence, flowering intensity, number of fruits, single fruit weight, red over color and russet frequency, reaching a maximum increase in predictive ability of 0.07 for flowering intensity. For red over color, the model A+A×E performed better than its counterpart, the model G+G×E, achieving a 0.01 higher predictive ability. Although a significant improvement in predictive ability was observed for model A+D+A×E+D×E for five traits compared to model G, the model A+D+A×E+D×E did not provide any additional improvement compared to simpler models A+D and A+A×E. For models G+W, G+W+G×W, and G+G×E+W+G×W, no additional improvement in predictive ability was found when compared with model G, except for model G+G×E+W+G×W, which showed a significantly higher predictive ability for flowering intensity, but this value was 0.01 lower than for the simpler model G+G×E.

**Figure 5:**
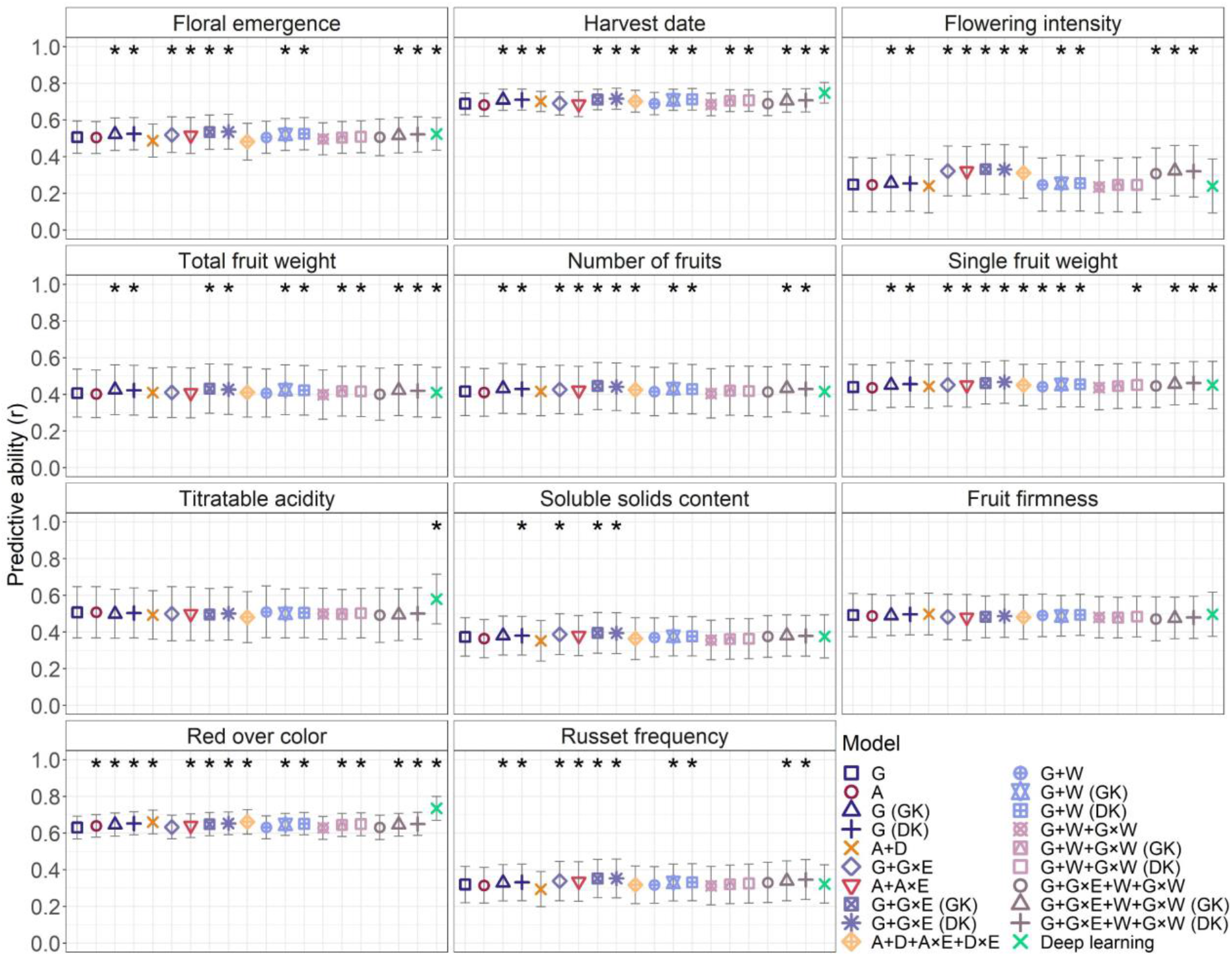
Average predictive abilities for eleven traits estimated using statistical and deep learning genomic prediction models. The statistical genomic prediction models were based on combinations of the genomic (G), additive (A), dominance (D), and enviromic (W) effects, interactions (×) with the vector of environments (E), and interactions between the genomic and environmental effects (G×W). The relationship matrices for the different effects in the statistical genomic prediction models were constructed using the G-BLUP approach or, where indicated, the Gaussian kernel (GK) or deep kernel (DK). For each model and trait, the predictive abilities were estimated separately for every environment nested in the folds and repetitions of the cross-validation and then averaged. Error bars correspond to standard deviation around the mean. Prior to averaging, the Wilcoxon signed-rank exact test was used to compare predictive abilities for each model with the predictive abilities obtained for the benchmark model G based on G-BLUP. Predictive abilities significantly greater than those of the benchmark model are labeled with an asterisk (Bonferroni-corrected α = 0.05).

The comparison between model G based on G-BLUP, model G (GK) employing the Gaussian kernel, and model G (DK) utilizing the Deep kernel revealed a significant improvement in predictive ability, ranging from 0.01 to 0.02 for most traits (Figure 5, Table S5). For nine out of eleven traits, both models G+G×E (GK) and G+G×E (DK) exhibited a significant enhancement in predictive ability compared to model G. Additionally, these models demonstrated an increased predictive ability ranging from 0.01 to 0.03 when compared to the G+G×E model based on G-BLUP. Specifically, for flowering intensity, models G+G×E (GK) and G+G×E (DK) displayed a 0.08 higher predictive ability than model G. The models G+W (GK) and G+W (DK) resulted in a significant increase in predictive ability of 0.01 to 0.02 across eight traits compared to the model G. However, these models did not surpass the predictive ability of the models G (GK) and G (DK). The inclusion of additional effects resulting in models G+W+G×W (GK), G+W+G×W (DK), G+G×E+W+G×W (GK), and G+G×E+W+G×W (DK) did not improve predictive ability compared to models with fewer components based on G-BLUP, Gaussian kernel and Deep kernel.

The deep learning genomic prediction model demonstrated significantly higher predictive abilities than model G for six out of the eleven traits studied (Figure 5, Table S5). For harvest date, titratable acidity and red over color, the deep learning genomic prediction model outperformed all statistical genomic prediction models tested. The increase in predictive ability compared to model G was 0.06 for harvest date, 0.07 for titratable acidity, and 0.10 for red over color.

### Model efficiency

Averaged across all traits, the predictive ability ranged from 0.45 to 0.49 for the compared models (Figure 6, Table S6). Based on these average predictive abilities, the model G+W+G×W emerged as the least efficient. Conversely, the models G+G×E (GK) and G+G×E (DK) showed the highest average predictive abilities, followed by the deep learning genomic prediction model with the average predictive ability of 0.48. Among the compared statistical and deep learning genomic prediction models, the average computational time required for the creation of the relationship matrices and model structures to the completion of the model fitting varied from 2 to 93 minutes (Figure 6, Table S6). The longest average computational time of 93 minutes was observed for the most complex model G+G×E+W+G×W (GK). The shortest average computational time among all models of 2 minutes was obtained when the analyses using the deep learning genomic prediction model were performed on a graphics processing unit (GPU). In contrast, longer average computational time of 42 minutes resulted from running the deep learning genomic prediction model on a central processing unit (CPU), which was also used for the statistical genomic prediction models. Among the statistical genomic prediction models, the benchmark model G based on G-BLUP was the fastest with an average computational time of 6 minutes. Comparing the average computational time with the average predictive ability across all traits studied, the G+G×E (DK) model stood out as the most efficient in terms of predictive ability, with a computational time of 10 minutes, which was only 8 and 4 minutes longer than the fastest deep learning and statistical genomic prediction models, respectively (Figure 6, Table S6).

**Figure 6:**
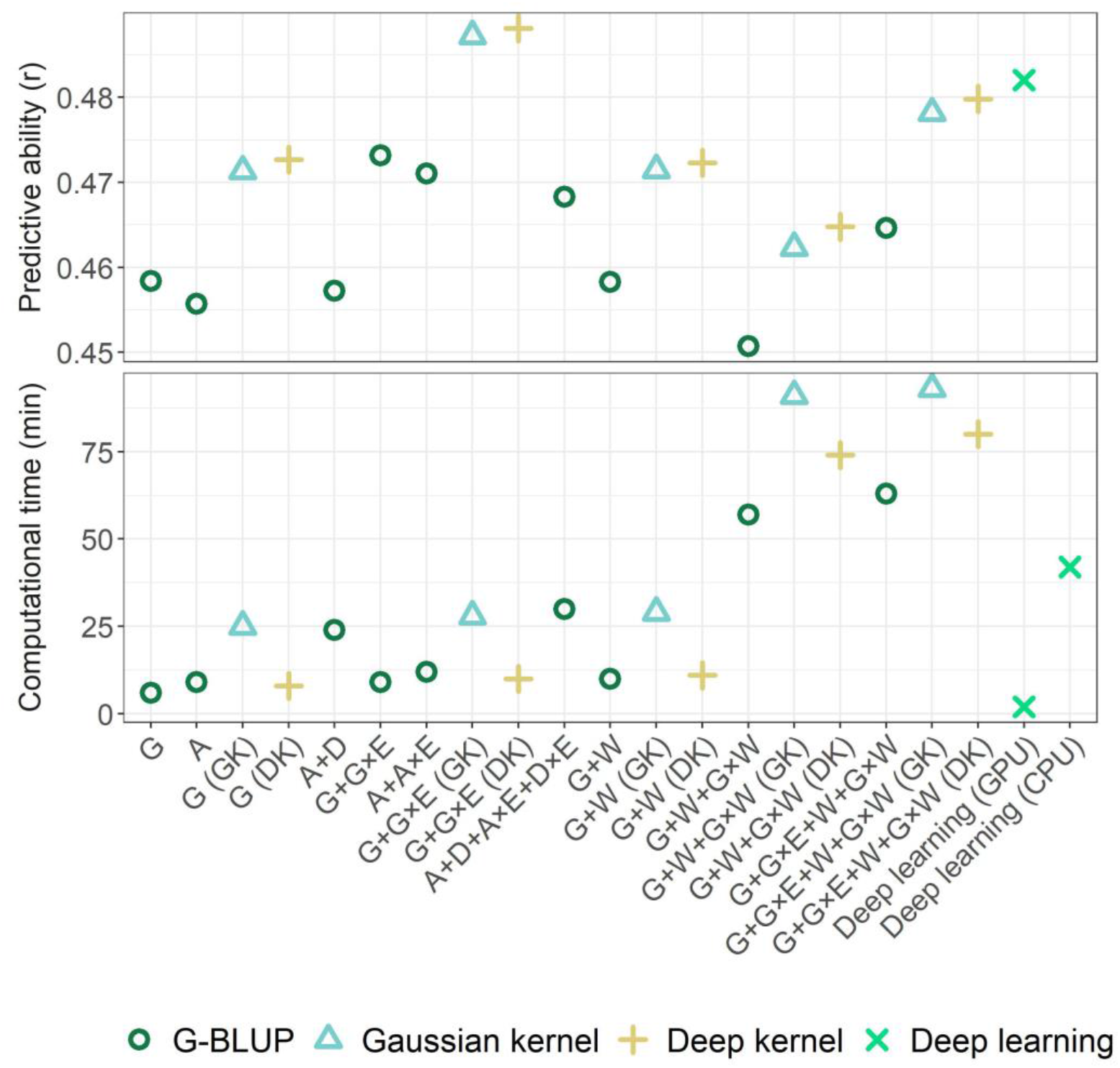
Comparison of predictive ability and computational time averaged across all studied traits. The statistical genomic prediction models were based on combinations of the genomic (G), additive (A), dominance (D), and enviromic (W) effects, interactions (×) with the vector of environments (E), and interactions between the genomic and environmental effects (G×W). The relationship matrices for the different effects in the statistical genomic prediction models were constructed using the G-BLUP approach or, where indicated, the Gaussian kernel (GK) or deep kernel (DK). To compare computational times, GPU-based deep learning computations were partially replicated on the CPU, which was used for statistical genomic prediction models (CPU – central processing unit, GPU – graphics processing unit). CPU was not used to assess predictive ability of the deep learning genomic prediction model.

## DISCUSSION

This study provides insights into the complexities of multi-environmental genomic prediction in quantitative apple traits. The incorporation of different sources of variation in the form of model components, and the comparison of predictive abilities and computational times between statistical genomic prediction models and a deep learning approach contribute to advancing the understanding of efficient genomic prediction methodologies. The findings highlight the need for a nuanced approach, considering the specific traits and modelling approaches in plant breeding applications.

### Modelling genotype-by-environment interaction

In the context of genomic prediction across environments (defined as combinations of location and year), this work underscored a significant improvement in predictive ability when employing genomic prediction models based on G-BLUP that integrate both main marker effects and the interaction effects of markers and environments, as it has been described by previous studies (Jarquí n et al., 2014; Jung et al., 2022; Lopez-Cruz et al., 2015). Compared to the benchmark genomic prediction model implementing exclusively the main marker effects, Jung et al. (2022) reported up to 0.07 increase in predictive ability for apple traits by integrating the random effects for G×E using the software package ‘BGLR’ (Pe rez & de los Campos, 2014). In this study deploying the newer software ‘BGGE’ (Granato et al., 2018), an analogous model comparison based on the same plant material but including two additional years of phenotypic data showed comparable improvements in predictive ability of up to 0.07. Average predictive ability across eleven studied traits for models incorporating G×E using G-BLUP was 0.01 lower compared to the average predictive ability for the same traits reported previously (Table S5, Jung et al., 2022). As the predictive ability of G×E models based on G-BLUP was similar in ‘BGLR’ and ‘BGGE’ (Granato et al., 2018), the difference in predictive ability was likely due to the changes in the phenotypic dataset between the compared studies.

### Additive and dominance effects

Orthogonal partitioning of variances implies that the proportions of additive genomic effects remain unchanged when additional effects, such as dominance, are introduced into the genomic prediction model (A lvarez-Castro & Carlborg, 2007). Despite using the natural and orthogonal interactions (NOIA) procedure for orthogonal partitioning of additive and dominant variances (A lvarez-Castro & Carlborg, 2007; Vitezica et al., 2017), our results indicate nonorthogonality when comparing models A and A+D. Specifically, the comparison of these models showed a 22% reduction in the average proportion of additive variance across all studied traits for model A+D, and a 1% decrease in the total average variance explained by model A+D (Table S2, Table S3). Similar results have been found in different crops, where the extension of models analogous to model A with dominance effects has often led to reduced estimates of additive variance components, and sometimes even to a reduction in the total explained variance below the level achieved by model A (Amadeu et al., 2020; Costa-Neto, Fritsche-Neto, et al., 2021; Roth et al., 2022; Yadav et al., 2021). While an earlier study showed that dominance variance was overestimated when inbreeding was not taken into account (Vitezica et al., 2018), our variance decomposition showed no signs of upwardly biased estimates of dominance variance in model A+D compared to A+D (inb) (Table S2). Our results likely point to potential problems in variance estimation caused by linkage disequilibrium (Roth et al., 2022; Vitezica et al., 2017), which is prevalent in breeding material such as that contained in the apple REFPOP. Beside the violation of the assumption of linkage equilibrium, the incorrect variance partitioning may have resulted from fitting multiple genetic and genotype-by-environment interaction effects within the context of multi-environmental genomic prediction, which deserves further investigation. A preliminary analysis outside the scope of this study indicated that orthogonality was restored when conducting analyses on across-location clonal values (results not shown). Nevertheless, compared to other approaches to modelling non-additive effects, the NOIA approach retains the advantage of allowing deviations from the Hardy-Weinberg equilibrium (Vitezica et al., 2017). Furthermore, the limitations in disentangling the additive and dominance effects had no negative impact on predictive ability of the model A+D compared to model A (Figure 6).

Model A+D provided improvements in predictive ability of 0.01 and 0.03 compared to model G for the traits harvest date and red over color, respectively (Table S5). Harvest date and red over color exhibited a distinctive oligogenic architecture among the studied traits (Jung et al., 2022), potentially accounting for the favorable influence of incorporating dominance effects in genomic prediction. Compared to more complex traits, the dominance effects of genes with large effects on traits may be easier to capture. For harvest date, a predominantly co-dominant mode of inheritance was proposed for the *NAC18.1* locus, which is a principal regulator of ripening traits in apple (Watts et al., 2023). For red over color, earlier literature have suggested that anthocyanin synthesis in apple skin was determined by a major dominant gene (Cheng et al., 1996; Takos et al., 2006), which was designated as *MdMYB1* (Takos et al., 2006). A semi-dominant effect of the allele *MdMYB1-1* associated with red fruit skin color was later reported by Moriya et al. (2017). Additionally, our results indicated the possibility of dominance effects for single fruit weight, a trait that showed an increase in predictive ability of 0.01 using model A+D (inb) compared to model A+D (Table S5). The applied deep learning genomic prediction model pointed to a strong association with single fruit weight for the marker AX-115591267 on chromosome 1 at 28.6 Mb, which showed the highest absolute mean SHAP for this trait (Figure S5). This marker was located in a region previously associated with single fruit weight at 27.8 Mb (Jung et al., 2022), and a comparison of phenotypic values for this marker indicated dominance effects acting at this locus (Figure S6).

As suggested by the visually perceived similarity of the standard genomic and additive genomic relationship matrices based on G-BLUP, the models G and A as well as G+G×E and A+A×E were mostly equivalent in their respective predictive ability. Despite the potential for improved predictive ability from the implementation of orthogonal additive and dominance coefficients for the construction of genomic relationship matrices using G-BLUP and their combination with the fixed effect of inbreeding (Roth et al., 2022; Yadav et al., 2021), limited improvement in predictive ability was observed in this study for the models A+D and A+D+A×E+D×E as well as for the model structures implementing inbreeding (Table S5). These results are consistent with previous observation of nearly identical predictive ability between genomic prediction models with and without dominance effects analyzing quantitative traits in perennial fruit species such as blueberry (Amadeu et al., 2020).

### Non-genetic effects from envirotyping

As suggested by moderate differences in daily weather variables among years and locations, and the low differentiation between environments within a location in the enviromic relationship matrix, environmental covariates discriminated well between locations but weakly between specific environments. This could likely be explained by the larger number of soil covariates (22) than weather covariates (6), and the lack of variability between years for the soil covariates due to their single measurement at each orchard location in 2016. Additionally, the precipitation variable, which could have aided in distinguishing between environments, had to be excluded from the analysis. This decision was prompted by the confounding of precipitation with irrigation at some apple REFPOP locations. Nevertheless, the enviromic-based effects explained a substantial part of the phenotypic variance, especially for floral emergence known to be strongly affected by the environment (Jung et al., 2022). Although a large proportion of phenotypic variance was explained here by the enviromic-based effects, and these effects have been shown to positively influence predictive ability in other crops (Costa-Neto, Fritsche-Neto, et al., 2021; Jarquí n et al., 2014), they have not resulted in any increase in predictive ability for apple traits. For productivity traits such as flowering intensity, which depends on flower bud formation during the previous vegetation season, the models could likely benefit from including prior-year environmental data in the construction of the enviromic matrix.

### Alternative kernels

Similar to previous reports that have shown increased predictive ability when Gaussian kernel and Deep kernel were applied (Costa-Neto, Fritsche-Neto, et al., 2021; Cuevas et al., 2019), these kernels resulted in a modest but significant improvement in predictive ability of 0.01–0.02 for most of the studied traits. The Gaussian kernel proved particularly suitable for capturing variance attributed to G×E, albeit with a longer computational time compared to the Deep kernel. Model structures based on the Deep kernel generally explained a smaller proportion of phenotypic variance than those using the Gaussian kernel and G-BLUP. This characteristic rendered Deep kernel less suitable for evaluating trait genetic architecture. Nevertheless, the Deep kernel-based models demonstrated improved predictive abilities, equivalent to those of Gaussian kernel-based models. Overall, both alternative kernels proved to be efficient substitutes for G-BLUP.

### Deep learning for genomic prediction

Specifically for each trait and cross-validation fold, the dimensional reduction of the marker dataset to a subset of 1,000 SNPs selected by a gradient boosting algorithm, extended with known marker-trait associations, allowed a time-efficient implementation of a deep learning approach for multi-environmental genomic prediction in apple. The studied deep learning approach combined feature streams derived from marker information with streams incorporating weather and soil variables. It resulted in stream contributions that effectively represented trait genetic architectures described in this and previous studies using statistical genomic prediction models (Jung et al., 2022).

The trait genetic architecture for harvest date was particularly well captured, with a 72% contribution from the SNP stream. Harvest date was previously described as oligogenic trait with significant large-effect marker associations found on chromosomes 3, 10 and 16 using the apple REFPOP dataset (Jung et al., 2020, 2022). The strongest of these associations on chromosome 3 at 30.7 Mb (Jung et al., 2022) was located in a major locus *NAC18.1* associated with harvest date and multiple ripening traits (Migicovsky et al., 2016; Watts et al., 2023). The deep learning genomic prediction model proved efficient in capturing this major locus, as the three SNPs with the highest absolute mean SHAP were located on chromosome 3 at 29.2, 30.7, and 30.8 Mb, the marker AX-115366114 at 30.7 Mb being strongly associated with harvest date according to our previous study (Jung et al., 2022). Moreover, the deep learning genomic prediction model outperformed the benchmark statistical genomic prediction model G for harvest date, significantly improving predictive ability by 0.06 and achieving the highest predictive ability among all tested models at 0.75.

Red over color has shown similar predictive ability and trait genetic architecture as harvest date in this and previous studies based on statistical genomic prediction models (Jung et al., 2022). The SNPs associated with *MdMYB1* transcription factor on chromosome 9, which regulates red pigmentation of apple skin (Takos et al., 2006), translated into large absolute mean SHAP values and predictive ability improved by 0.10 compared to model G. Similar results were observed for titratable acidity, where large absolute mean SHAP were found, and the three SNPs with the largest SHAP were located on chromosome 8 at 10.7, 10.8, and 11.8 Mb. Two large-effect loci are known for acidity in apple, namely *Ma* on chromosome 16 and *Ma3* on chromosome 8 (Verma et al., 2019). The SNPs on chromosome 8 indicated a strong association with the *Ma3* locus, and they colocalized with the SNP marker predictive for this locus at 10.9 Mb (Rymenants et al., 2020). The maximum relative SHAP contribution for the SNP stream of 80% was reached for titratable acidity. Moreover, the predictive ability of the deep learning genomic prediction model for titratable acidity was improved by 0.07 compared to the statistical genomic prediction model G. Our results for harvest date, red over color, and titratable acidity showed that high relative and absolute SHAP values can serve as predictors of improved deep learning genomic prediction model performance, and that the applied deep learning approach can precisely predict apple traits characterized by oligogenic architecture.

Unless very large datasets are examined, the predictive ability of deep learning approaches typically falls below that of conventional models for genomic prediction (Montesinos-Lo pez et al., 2021). Between the three traits with predictive ability superior to all other compared statistical genomic prediction models, the sizes of datasets showed large differences (total number of training instances of 12,428 for harvest date, 10,317 for red over color, and 2,879 for titratable acidity, ranging from 2,879 to 12,461 across all traits, Table S1). Although the number of available environments ranged from the minimum of eight for titratable acidity to the maximum of 25 for harvest date, similar improvement in predictive ability was reached for these traits using the applied deep learning approach. As the improvements in predictive ability for harvest date, red over color, and titratable acidity were observed independently from the number of training instances, the size of the phenotypic dataset is unlikely to have affected our predictions.

### Multi-environmental genomic selection in apple breeding

The establishment of multi-environmental genomic selection in apple has been constrained by several factors, including the costly collection of extensive multi-environmental datasets and computational limitations. The phenotyping efforts in the apple REFPOP yielded an unprecedented dataset in terms of trait-environment combinations (Jung et al., 2022), which has been expanded in this study with two additional years of phenotyping. This dataset now encompasses phenotypic data for eleven traits across up to 25 environments. The availability of this dataset has enabled the implementation of multi-environmental genomic prediction models within a computationally efficient framework, laying the groundwork for the practical application of multi-environmental genomic selection in apple.

The approach to multi-environmental genomic prediction of apple traits used in this study diverges from the traditional understanding of environments in apple tree cultivation. In practice, apple trees remain stationary in the same location across multiple years. This stationary nature of apple cultivation implies that the effects of yearly climatic variations are superimposed on the same geographical location, whereas the genomic prediction approach treats each year-location combination as a distinct environment. Nevertheless, breeding values for apple genotypes lacking phenotypic information can be predicted across diverse environmental conditions using the genomic prediction models trained in this study.

Among all predictions obtained, the model G+G×E applying Gaussian and Deep kernels improved predictive abilities for most traits (except for titratable acidity and fruit firmness, where it showed results comparable to those of the benchmark model G). Therefore, the model G+G×E proved to be a universally effective solution for multi-environmental genomic prediction in the studied apple traits. Additionally, the G+G×E model, along with other statistical genomic prediction models tested, was outperformed by the applied deep learning approach for three traits with simpler genetic architectures (harvest date, titratable acidity, and red over color). Depending on the genetic architecture of the trait, either the G+G×E model or the deep learning approach can be recommended for multi-environmental genomic predictions, leading to informed breeding decisions, and assisting in the selection of cultivars more adaptable to future climates.

## MATERIALS AND METHODS

### Plant material

The apple REFPOP comprised 265 progenies from 27 biparental families generated by European breeding programs, along with 269 diverse accessions (Jung et al., 2020). This study focused on five locations: (i) Rillaar, Belgium, (ii) Angers, France, (iii) Laimburg, Italy, (iv) Lleida, Spain, and (v) Waedenswil, Switzerland. At each location, all genotypes were generally represented by two trees and planted in 2016 using a randomized complete block design. Three control genotypes, namely ’Gala’, ’Golden Delicious’, and ’CIVG198’, were replicated up to 22 times at each location. The cultivation followed the common agricultural practices specific to each location, incorporating integrated plant protection methods.

### Phenotyping

Phenotyping of the eleven traits followed the methodology described by Jung et al. (2022). Individual trees, representing genotype replicates, were used for trait measurement. Floral emergence was determined in Julian days, marking the date when the first 10% of flowers opened. Flowering intensity was evaluated on a nine-grade scale, indicating the percentage of existing flowers relative to the maximum potential number of flowers. Fruits were harvested on harvest dates, determined in Julian days, based on expert estimates of fruit ripening. Total fruit weight (kg) and fruit number were recorded to assess production per tree. Single fruit weight (g) was estimated by dividing the total fruit weight by the number of fruits. Titratable acidity (g/l), soluble solids content (°Brix), and fruit firmness (g/cm^2^) were measured within one week post-harvest using an automated instrument Pimprenelle (Setop, France). Red over color, representing the percentage of red fruit skin, was assessed on a six-grade scale. Russet frequency indicated the percentage of fruits exhibiting russet skin. Further information regarding the evaluation of the eleven traits is available in Jung et al. (2022). For the different traits, the assessment spanned a period of up to five years from 2018 to 2022 and was performed at up to five locations.

### Envirotyping

Hourly measurements of temperature (°C) at 2 m above soil level, relative humidity (%) and global radiation (W/m2) were obtained from the weather stations near the apple REFPOP orchards from 2018 to 2022. Precipitation (mm) was not taken into consideration in this study due to irrigation practices in part of the orchard locations.

In each apple REFPOP orchard between May 12 and June 9, 2016, a total of six soil samples were collected from three distinct sampling points and two soil depths (approximately 1–20 cm and 20–40 cm). In the accredited Laboratory for Soil and Plant Analysis of Laimburg Research Centre, Italy, the soil samples were analyzed for (i) organic carbon (% humus), (ii) pH, (iii) carbonate test, expressed as low to medium, high, very high or no carbonate content, (iv) carbonate requirement (dt/ha CaO), (v) phosphorus (mg/100 g P_2_O_5_), (vi) potassium (mg/100 g K_2_O), (vii) magnesium (mg/100 g), (viii) boron (mg/kg), (ix) manganese (mg/kg), (x) copper (mg/kg), and (xi) zinc (mg/kg).

### Genotyping

As detailed by Jung et al. (2020), the apple REFPOP underwent genotyping for biallelic single nucleotide polymorphisms (SNPs) through a dual approach utilizing the Illumina Infinium® 20K SNP genotyping array (Bianco et al., 2014) and the Affymetrix Axiom® Apple 480K SNP genotyping array (Bianco et al., 2016). By employing the Beagle 4.0 software (Browning & Browning, 2007) and incorporating pedigree information (Muranty et al., 2020), the obtained SNP sets were integrated through imputation, ultimately yielding a genomic dataset of 303,239 biallelic SNPs. All SNP positions were based on the doubled haploid GDDH13 (v1.1) reference genome (Daccord et al., 2017).

### Phenotypic data preprocessing

Analyses of phenotypic data were conducted to ensure high data quality by addressing low heritability, spatial heterogeneity, and eliminating outliers. The statistical model for the phenotypic data preprocessing was fitted via restricted maximum likelihood using the R package ‘lme4’ (v.1.1-28) (Bates et al., 2015) as:

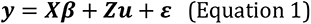

where *y* was the vector of the response variable, *X* the design matrix for the fixed effects, *β* the vector of the fixed effects, *Z* was the design matrix for the random effects, *u* the vector of the random effects assuming *µ*∼*N*(0, ∑) with ∑ being the variance–covariance matrix of the random effects and ε the vector of the random errors assuming *ε*∼*N*(0, σ_ε_^2^*I*) with σ_ε_^2^ being the error variance and *I* the identity matrix.

Separately for each trait and environment (combined factor of location and year), raw phenotypic values for each genotype replicate (total fruit weight and fruit number were log-transformed) were used as response variable to fit a random-effects model with a random effect of genotype following the Equation 1. From the variance components of the random-effects model, the environment-specific clonal mean heritability was calculated as:

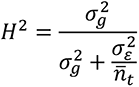

where σ_g_^2^ was the genotypic variance and *n̄*_*t*_ the mean number of genotype replications. The environment-specific clonal mean heritability was used to remove trait-environment combinations with the heritability value below 0.1.

To account for spatial variation in the orchards, spatial heterogeneity in the raw phenotypic data was modeled separately for each trait-environment combination using the spatial analysis of field trials with splines (‘SpATS’ (v.1.0-11)) (Rodrí guez-A lvarez et al., 2018) as described by Jung et al. (2020). From the fitted SpATS objects, the adjusted phenotypic values of each genotype and the adjusted phenotypic values of each tree were obtained.

The adjusted phenotypic values of each genotype were used as response variable for fitting a mixed-effects model with a fixed effect of environment and a random effect of genotype following Equation 1. Subsequently, the outliers were detected using Bonferroni–Holm test to judge residuals standardized by the re-scaled median absolute deviation (BH-MADR) as described by Bernal-Vasquez et al. (2016). The identified outliers were removed and the remaining trait- and environment-specific adjusted phenotypic values of each genotype were further denoted as adjusted means. The adjusted means for the eleven studied traits were compared separately for each year and location using the pairwise Pearson’s correlations and significance tests implemented in the R package ‘corrplot’ (v.0.92) (Wei & Simko, 2021). The significance levels of 0.05, 0.01, and 0.001 were Bonferroni-corrected by dividing them by the total number of pairwise comparisons among the eleven traits.

Following Equation 1, the adjusted phenotypic values of each tree served as the response variable in fitting a mixed-effects model, denoted here as the phenotypic model. This model included the fixed effects of environment (E), the random effects of genotype (g), and random effects of genotype-by-environment interaction (g×E). The proportions of phenotypic variance explained by the random effects were extracted from the model fit for comparison with the statistical genomic prediction models.

### Enviromic data preprocessing

The enviromic data were restructured to acquire appropriate inputs for the subsequent modelling. Daily temperature means, daily humidity means, and daily radiation sums were calculated from the hourly measurements. These three daily weather variables were visualized applying local regression curves estimated using Loess with a span of 0.1.

Inspired by Jarquí n et al. (2014), the three daily weather variables were processed to create six environmental covariates by dividing each growing season into two periods based on crop phenology. The two periods were defined separately for each environment. The first period extended for 80 days, concluding on the day when 90% of the genotypes flowered, determined from adjusted means for floral emergence. The second period followed the first until the day when 90% of the genotypes were harvested, as indicated by the adjusted means for harvest date. Different approaches to defining the first period were employed for two environments where adjusted means for floral emergence were unavailable. In the case of the environment ESP.2020, which was excluded due to low heritability, the adjusted phenotypic values of each genotype were used to estimate the day when 90% of the genotypes flowered. For ESP.2022, where floral emergence scores were missing, the end date of the first period was estimated based on varieties cultivated near the apple REFPOP. Daily temperature means, daily humidity means, and daily radiation sums were summed over each respective period, resulting in six environmental covariates. Additionally, 22 environmental covariates were obtained as mean values of eleven soil characteristics calculated per location and the level of soil depth. All 28 environmental covariates were collected in the *q* × *z* matrix of environmental covariates *M*_*W*_ with *q* environments and *z* environmental covariates, which was then scaled and centered to the mean of zero and standard deviation of one.

### Marker matrices

Three marker matrices were constructed based on the genomic dataset of biallelic SNPs. The first matrix followed the standard allele coding, where a SNP was assigned the value 0 when the individual (i.e., genotype) was homozygous for the first allele (*a*), 1 when the genotype was heterozygous, and 2 when the genotype was homozygous for the second allele (*A*). The allele coding can be referred to as coefficients in the marker matrix. Therefore, the *n* × *m* marker matrix of the standard coefficients *M*_*G*_ with *i* = 1, … , *n* genotypes and *j* = 1, … , *m* markers was:

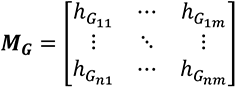

where the element ℎ_*Gij*_for the *i*th genotype and *j*th marker was equal to:

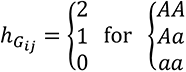

with *AA*, *Aa* and *aa* being the combinations of the alleles *a* and *A* at the marker *j*. Each column of the matrix *M*_*G*_ was scaled and centered to the mean of zero and standard deviation of one.

The second and third marker matrices followed the natural and orthogonal interactions (NOIA) model (A lvarez-Castro & Carlborg, 2007) as implemented by Vitezica et al. (2017). These matrices were estimated from the elements of the marker matrix of the standard coefficients *M*_*G*_ and had the same dimension. The element ℎ_*Aij*_ for the *n* × *m* marker matrix of additive coefficients *M*_*A*_ and the element *h*_*D_ij_*_ for the *n* × *m* marker matrix of dominance coefficients *M*_*D*_ were calculated as follows:

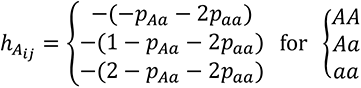

and

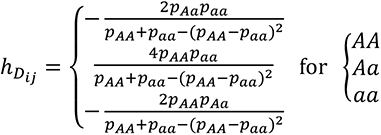

with *p*_*AA*_, *p*_*Aa*_ and *p*_*aa*_ being the relative frequencies for the allelic combinations *AA*, *Aa* and *aa* at marker *j*.

### Relationship matrices

The marker matrices *M*_*G*_, *M*_*A*_ and *M*_*D*_ and the matrix of environmental covariates *M*_*W*_ were used to estimate the standard genomic relationship matrix *K*_*G*_, the additive genomic relationship matrix *K*_*A*_, the dominance genomic relationship matrix *K*_*D*_, and the enviromic relationship matrix *K*_*W*_, respectively. Initially, all relationship matrices were created based on the genomic best linear unbiased predictor (G-BLUP) approach described by VanRaden (2008). The covariance matrix following the G-BLUP approach was obtained as:

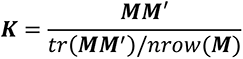

where *K* was a generic representation of the relationship matrix (*K*_*G*_, *K*_*A*_, *K*_*D*_ and *K*_*W*_), *M* was a generic representation of the marker matrices *M*_*G*_, *M*_*A*_ and *M*_*D*_ as well as the matrix of environmental covariates *M*_*W*_, and *nrow* was the number of genotypes for *M*_*G*_, *M*_*A*_ and *M*_*D*_ or the number of environments for *M*_*W*_.

Subsequently, two covariance matrix types, namely the Gaussian kernel (Gonza lez-Camacho et al., 2012) and Deep kernel (Cuevas et al., 2019), were examined as alternatives to the G-BLUP approach. The Gaussian kernel is a nonlinear method based on a bandwidth parameter that controls the decay rate of covariance between genotypes, and the percentile of the square of the Euclidean distance, which is a metric reflecting the genetic distance between genotypes. The Deep kernel is characterized by a nonlinear arc-cosine function, and its covariance matrix is designed to mimic a deep-learning model featuring a single hidden layer with many neurons. Applying these alternative approaches, the standard genomic and enviromic relationship matrices based on Gaussian kernel (*K*_*G_GK_*_ and *K*_*W_GK_*_ ) and Deep kernel (*K*_*G_DK_*_ and *K*_*W_DK_*_) were created. The Gaussian kernel and Deep kernel were implemented following the estimation process as detailed by Costa-Neto et al. (2021).

### Statistical genomic prediction model structures

The relationship matrices were used to create linear mixed model structures for the statistical genomic prediction models. Following Costa-Neto et al. (2021), the generic model structure was defined as:

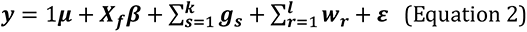

where *y* was the vector of the adjusted means for *n* genotypes across *q* environments, 1*µ* was the overall mean, *X*_*f*_ the design matrix for the fixed effects of environments, *β* the vector of the fixed effects, *g*_*s*_ the random vector for *s* = 1, … , *k* marker-based effects, *w*_*r*_ the random vector for *r* = 1, … , *l* enviromic-based effects, and *ε* the vector of the random errors assuming *ε*∼*N*(0, σ*_ε_*^2^*I*) with σ*_ε_*^2^being the error variance and *I* the identity matrix. All model structures were based on the G-BLUP approach to estimating the relationship matrices. When the alternatives to the G-BLUP were used, the model structures were additionally labeled with ‘(GK)’ for the Gaussian kernel and ‘(DK)’ for the Deep kernel. For all three approaches to estimating the relationship matrices, the function get_kernel of the R package ‘EnvRtype’ (v.1.1.1) (Costa-Neto, Galli, et al., 2021) was used to obtain the relationship matrices for genomic prediction.

### Models G, A, and A+D (main genotypic effects (MM))

Following the Equation 2, the model MM accounted for the marker-based effects (∑*_s_*_=1_^*k*^ *g*_*s*_ ≠ 0) without applying the enviromic-based effects (∑*_r=1_*^*l*^ *w*_*r*_ = 0). The *g*_*s*_ incorporated relationship matrices *K*_*G*_ (alternatively *K*_*G_GK_*_ or *K*_*G_GK_*_ *K*_*A*_ and *K*_*D*_ representing genomic (G), additive (A) and dominance (D) effects, respectively. These effects were applied individually or in combinations, resulting in model structures denoted as G (alternatively G (GK) and G (DK)), A and A+D.

### Models G+G×E, A+A×E, and A+D+A×E+D×E (single variance genotype × environment deviation (MDs))

Analogous to the model MM, the model MDs assumed ∑*_s_*_=1_^*k*^ *g*_*s*_ ≠ 0 and ∑*_l_*_=1_^*k*^*w*_*r*_ = 0 (Equation 2). In addition to the main effects G, A and D, the random interaction effects (×) with the vector of environments (E) were included, namely the G×E, A×E and D×E. This resulted in model structures G+G×E (alternatively G+G×E (GK) and G+G×E (DK)), A+A×E and A+D+A×E+D×E.

### Model G+W (enviromic-enriched MM (EMM))

The model EMM applied both the marker-based effects (∑*_s_*_=1_^*k*^ *g*_*s*_ ≠ 0) and the enviromic-based effects (∑*_r=1_*^*l*^ *w*_*r*_ ≠ 0) (Equation 2). Included were the main effects G and the enviromic effects (W), the latter being derived through the integration of the relationship matrix *K*_*W*_ (alternatively *K*_*W_GK_*_ and *K*_*W_DK_*_ ). The resulting model structure was G+W (alternatively G+W (GK) and G+W (DK)).

### Model G+W+G×W (reaction-norm MM (RNMM))

Building upon the model EMM, the model RNMM (∑*_s_*_=1_^*k*^ *g*_*s*_ ≠ 0 and ∑*_r_*_=1_^*l*^ *w*_*r*_ ≠ 0, Equation 2) extended the enviromic-based effects with a random interaction effect G×W. The obtained model structure was G+W+G×W (alternatively G+W+G×W (GK) and G+W+G×W (DK)).

### Model G+G×E+W+G×W (reaction-norm MDs (RNMDs))

The last of the compared models, the model RNMDs (∑*_s_*_=1_^*k*^ *g*_*s*_ ≠ 0 and ∑*_r_*_=1_^*l*^ *w*_*r*_ ≠ 0, Equation 2), combined the marker-based effects G and G×E with the enviromic-based effects W and G×W in a single model structure G+G×E+W+G×W (alternatively G+G×E+W+G×W (GK) and G+G×E+W+G×W (DK)).

### Fixed effect of inbreeding

The design matrix for the fixed effects *X*_*f*_ (Equation 2) was based on the vector of environments (E) for all model structures tested in this study. As described by previous authors (Roth et al., 2022; Vitezica et al., 2018), including an inbreeding coefficient as fixed effect can account for dominance effects and help to avoid overestimating the proportion of variance explained by the dominance model components. Hence, the model MM was additionally extended with the fixed effect of inbreeding contained in parameter *X*_*f*_, which was incorporated in the model structures denoted as A (inb) and A+D (inb). The inbreeding coefficient for each genotype was estimated from the marker matrix *M*_*G*_, calculated as the relative frequency of the homozygous allelic combinations *AA* and *aa* across all markers.

### Deep learning approach

The deep learning genomic prediction model was designed to be able to receive both genotypic and environmental data in the form of four streams. Genotypic data underwent feature selection in two different ways, generating input data for two different streams of the model: single nucleotide polymorphism (SNP) stream and principal component (PC) stream. First, to represent specific genetic variation, the most relevant 1,000 SNPs for each trait and fold were extracted from the marker matrix *M*_*G*_ with a gradient boosting regressor. The response variable for the gradient boosting model was derived from the means of the random effects of genotypes, which were extracted from a mixed-effects model. This mixed-effects model followed Equation 1, incorporating fixed effects of the environment (E) and random effects of genotype (g). Additionally, the SNPs associated with the studied traits as reported by Jung et al. (2022) were added to the existing pool of 1,000 SNPs within the SNP stream. Second, using the principal component analysis in related samples (PC-AiR) method (Kick et al., 2023), 58 principal components (PCs) capturing 100% of the genetic variation were extracted and used as input to represent the overall genetic variation. Daily weather variables and soil environmental covariates directly constituted the input for the weather and soil streams, respectively. The adjusted phenotypic means served as the response variables. All stream and response variables were scaled between -1 and 1. The model architecture was designed using ‘TensorFlow’ (v.2.10.0) and ‘Keras’ (v.2.10.0). All streams consisted of a variable number of dense layers except for the weather stream. In this case, the first layers were long short-term memory (LSTM), which excel at processing sequential data. The four streams processed the data independently and were concatenated after several layers. Further dense layers were placed before the output neuron to allow for data integration. For specific details on the model architecture, please refer to the provided GitHub link (https://github.com/MichaelaJung/Integrative-prediction). Models for each trait were trained and evaluated at different learning rates (1e^-4^, 1e^-5^, and 5e^-6^). When the training loss stopped improving, the training was stopped. The appropriate learning rate was decided for each trait based on the highest correlation and the lowest root mean squared error.

### Genomic prediction

All statistical and deep learning genomic prediction models were iteratively fitted in a five-fold cross-validation that was repeated five times, with genotypes being allocated to folds randomly and without replacement. The statistical genomic prediction model structures were solved using Bayesian hierarchical modeling implemented in the R package ‘BGGE’ (v.0.6.5) (Granato et al., 2018). The statistical genomic prediction models underwent 10,000 iterations of the Gibbs sampler, employing a thinning of 3 and discarding the initial 1,000 samples as burn-in.

### Relative contribution of model components

For the statistical genomic prediction models, each model fit from the cross-validation was used to obtain the proportions of variance explained by the various random effects. To explain the deep learning model predictions with respect to the inputs, the ‘GradientExplainer’ function from the ‘shap’ package (v.0.42.1) (Lundberg & Lee, 2017) was used to calculate approximated Shapley additive explanations (SHAP). It uses the gradients of the model to approximate SHAP values for each input, which estimates their contribution to the prediction. SHAP values were calculated for every instance of the test split in each repetition of the cross-validation. The absolute values of each feature were averaged to obtain absolute mean SHAP values. Furthermore, to investigate the contribution of each stream to the prediction, the absolute mean SHAP values were summed for every fold, and the relative SHAP contribution of each stream was obtained as a percentage.

### Assessment of predictive ability and computational time

For every statistical and deep learning genomic prediction model and trait, twenty-five estimates of predictive ability were generated for each environment, calculated as Pearson’s correlation coefficient between the adjusted means and predicted values. The Wilcoxon signed-rank exact test was used to compare predictive abilities of every model with the predictive abilities obtained for the benchmark model G with its model structure based on G-BLUP. From the resulting p-values, predictive abilities significantly greater than those of the benchmark model were identified using the Bonferroni-corrected significance threshold. The threshold was determined by dividing α = 0.05 by the total number of pairwise model comparisons among the eleven traits. For each cross-validation fold of the statistical genomic prediction models, the time was measured from the creation of the relationship matrices and model structures to the completion of the model fitting on central processing unit (CPU) nodes of the Euler cluster (ETH Zurich, Switzerland). The deep learning predictions were conducted on a graphics processing unit (GPU, NVIDIA GeForce RTX 3080), and their computational time was measured. To compare the computational times between statistical and deep learning genomic prediction models, deep learning computations were replicated only for the first five folds on the CPU. CPU was not used to assess predictive ability of the deep learning genomic prediction model.

All statistical analyses in this work were implemented in R (v.4.1.3) (R Core Team, 2022). The code for implementing, training, using, and explaining the deep learning genomic prediction models was written in Python (v.3.9.16).

## ACKNOWLEDGMENTS

The authors thank the field technicians and staff at INRAe IRHS and Experimental Unit (UE Horti), Angers, France, the Fruit Breeding Group at Agroscope in Waedenswil, Switzerland, and technical staff at all apple REFPOP sites for the maintenance of the orchards and phenotypic data collection. Phenotypic data collection was partially supported by the Horizon 2020 Framework Program of the European Union under grant agreement No 817970 (project INVITE: “Innovations in plant variety testing in Europe to foster the introduction of new varieties better adapted to varying biotic and abiotic conditions and to more sustainable crop management practices”). C.Q.-T. was supported by the European Union’s Horizon 2020 research and innovation program under the Marie Skłodowska-Curie grant agreement No 847585 – RESPONSE. This study was partially funded by the FOAG project “Apfelzukunft dank Zu chtung” (2020/17/AZZ).

## AUTHOR CONTRIBUTIONS

This research was conceived by M.J., M.Roth, M.J.A., W.G., F.L., H.M., B.S., and A.P. M.J., M.R., E.H., N.P., L.L., and F.D. contributed to data collection. M.J. carried out the statistical data analysis with the support of M.Roth., G.B., and A.P. C.Q.-T. performed the deep learning analysis in consultation with M.J., S.Y., and B.S. M.J. and C.Q.-T. wrote the article in consultation with M.Roth, M.J.A., H.M., M.R., W.G., F.L., S.Y., B.S., G.B., and A.P. All authors have read the manuscript and approved the version to be published.

## DATA AVAILABILITY

All SNP genotypic data used in this study have been deposited in the INRAe dataset archive at https://doi.org/10.15454/IOPGYF and https://doi.org/10.15454/1ERHGX. The raw phenotypic data are available in the INRAe dataset archive at https://doi.org/10.15454/VARJYJ. The code underlying this article is available in GitHub at https://github.com/MichaelaJung/Integrative-prediction. The phenotypic, enviromic, and imputed genomic data formatted as input files for the provided code are available in Zenodo at [to be updated upon article release].

