## Supplemental Figures for "Integrative multi-environmental genomic prediction in apple"

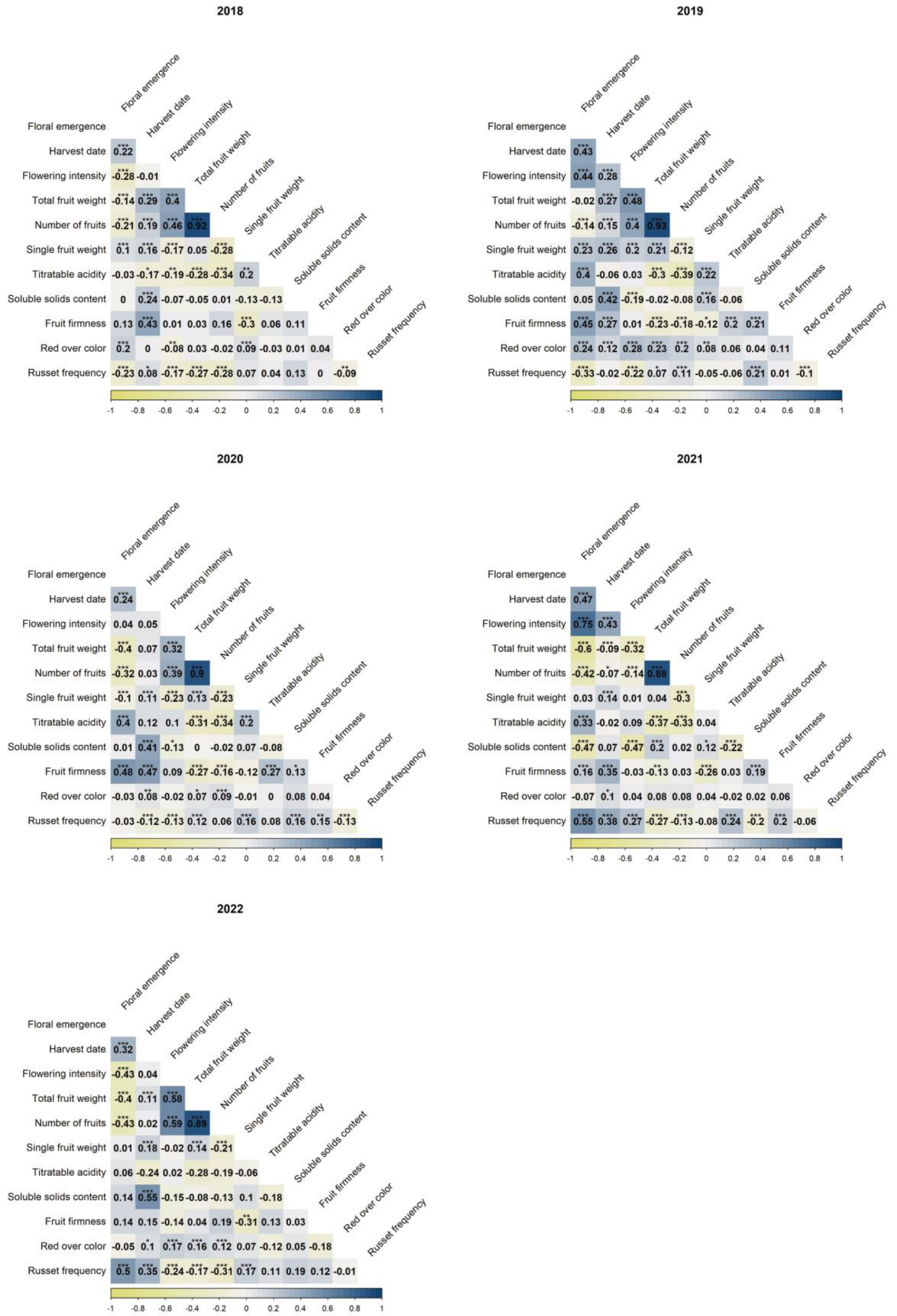


**Figure S1:** Pairwise Pearson's correlations of the adjusted means for eleven traits in five years (2018–2022) across locations. The asterisks indicate significant correlations at Bonferroni-corrected significance levels: * (α = 0.05), ** (α = 0.01), and *** (α = 0.001).

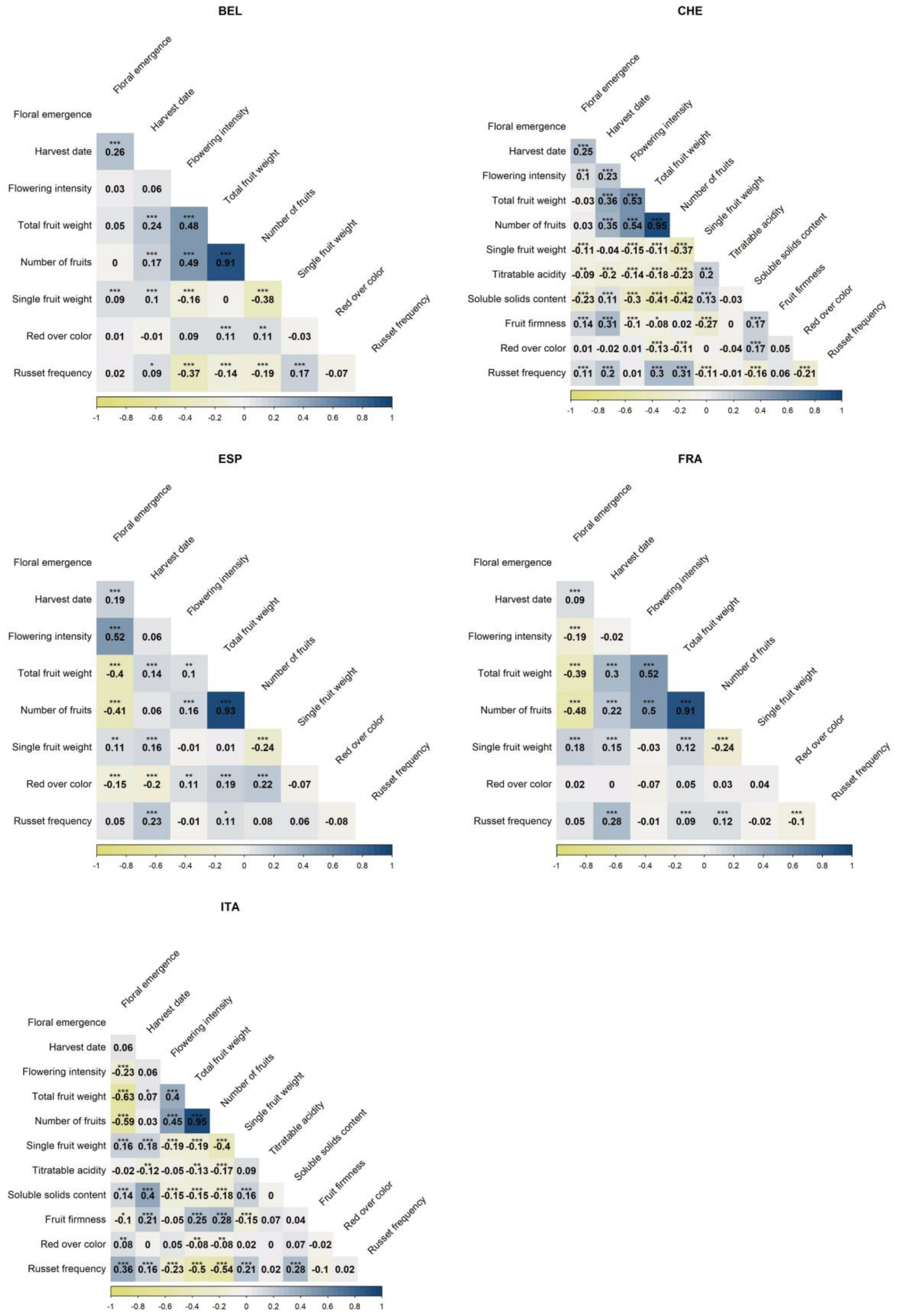


**Figure S2:** Pairwise Pearson's correlations of the adjusted means for eleven traits and five locations across years. The asterisks indicate significant correlations at Bonferroni-corrected significance levels: * (α = 0.05), ** (α = 0.01), and *** (α = 0.001). The locations correspond to Belgium (BEL), Switzerland (CHE), Spain (ESP), France (FRA) and Italy (ITA).

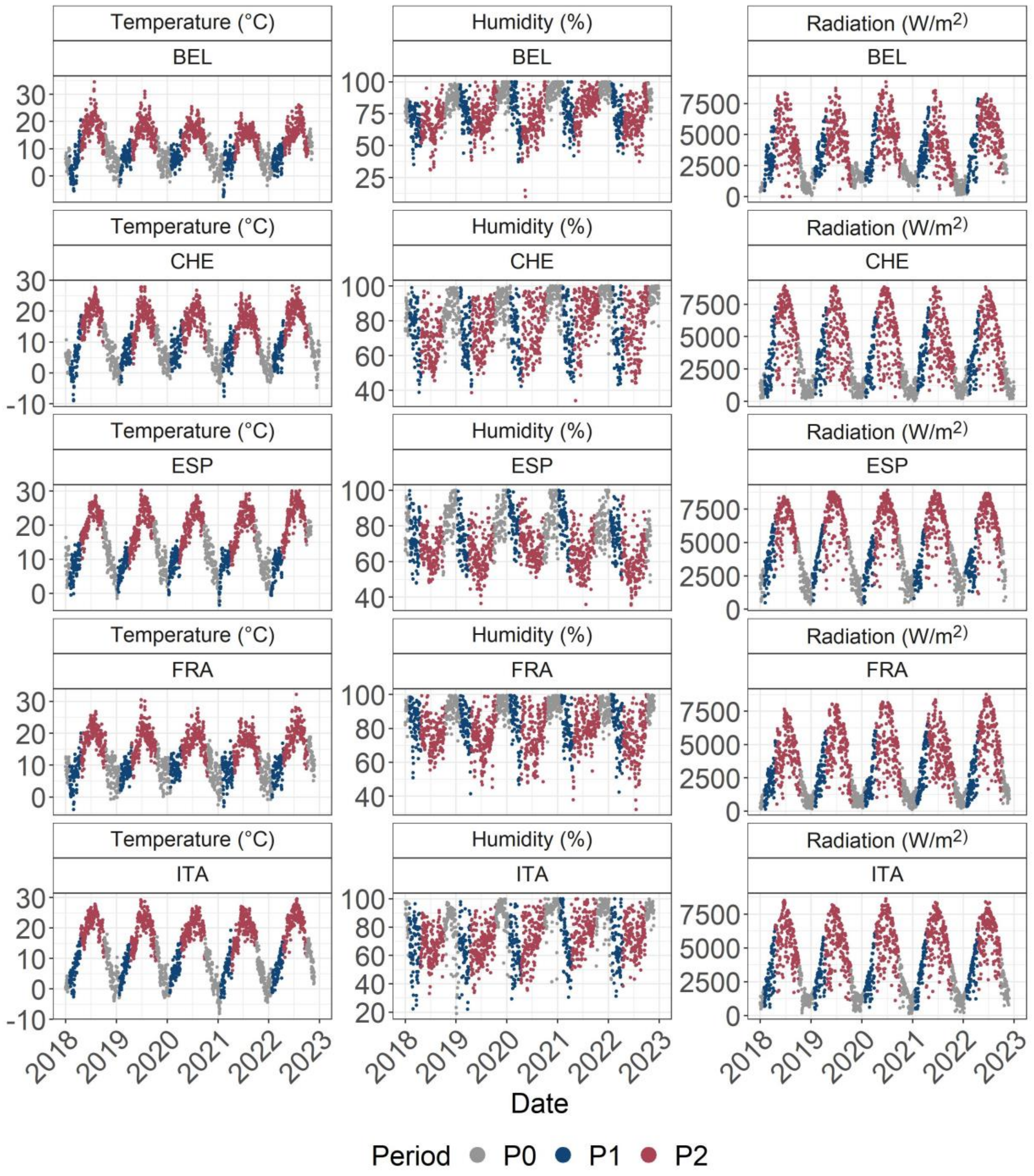


**Figure S3:** Daily temperature means, daily humidity means, and daily radiation sums for five locations and five years included in the study. The daily values are colored based on periods estimated from crop phenology. The first period (P1) extended for 80 days, concluding on the day when 90% of the genotypes flowered, determined from adjusted means for floral emergence. The second period (P2) followed the first until the day when 90% of the genotypes were harvested, as indicated by the adjusted means for harvest date. Daily values for P0 were excluded from the modelling process. The locations correspond to Belgium (BEL), Switzerland (CHE), Spain (ESP), France (FRA) and Italy (ITA).

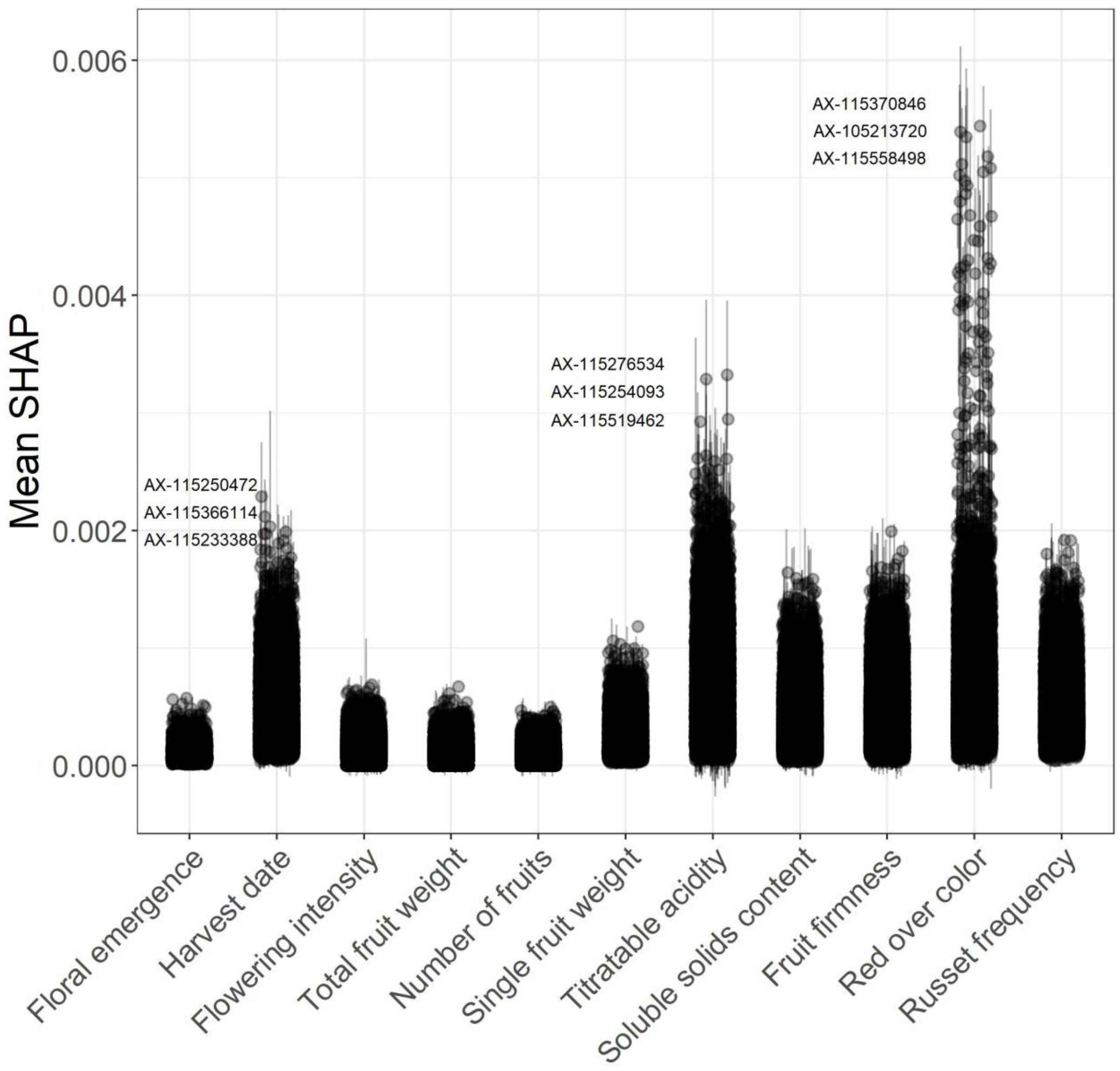


**Figure S4:** Absolute mean SHAP (Shapley additive explanations) for the SNP stream of the deep learning model. The labels (SNP numbers prefixed with AX) show names of the three most important SNPs for three traits with the largest absolute mean SHAP of the SNP stream. The error bars represent the standard deviation around the mean.

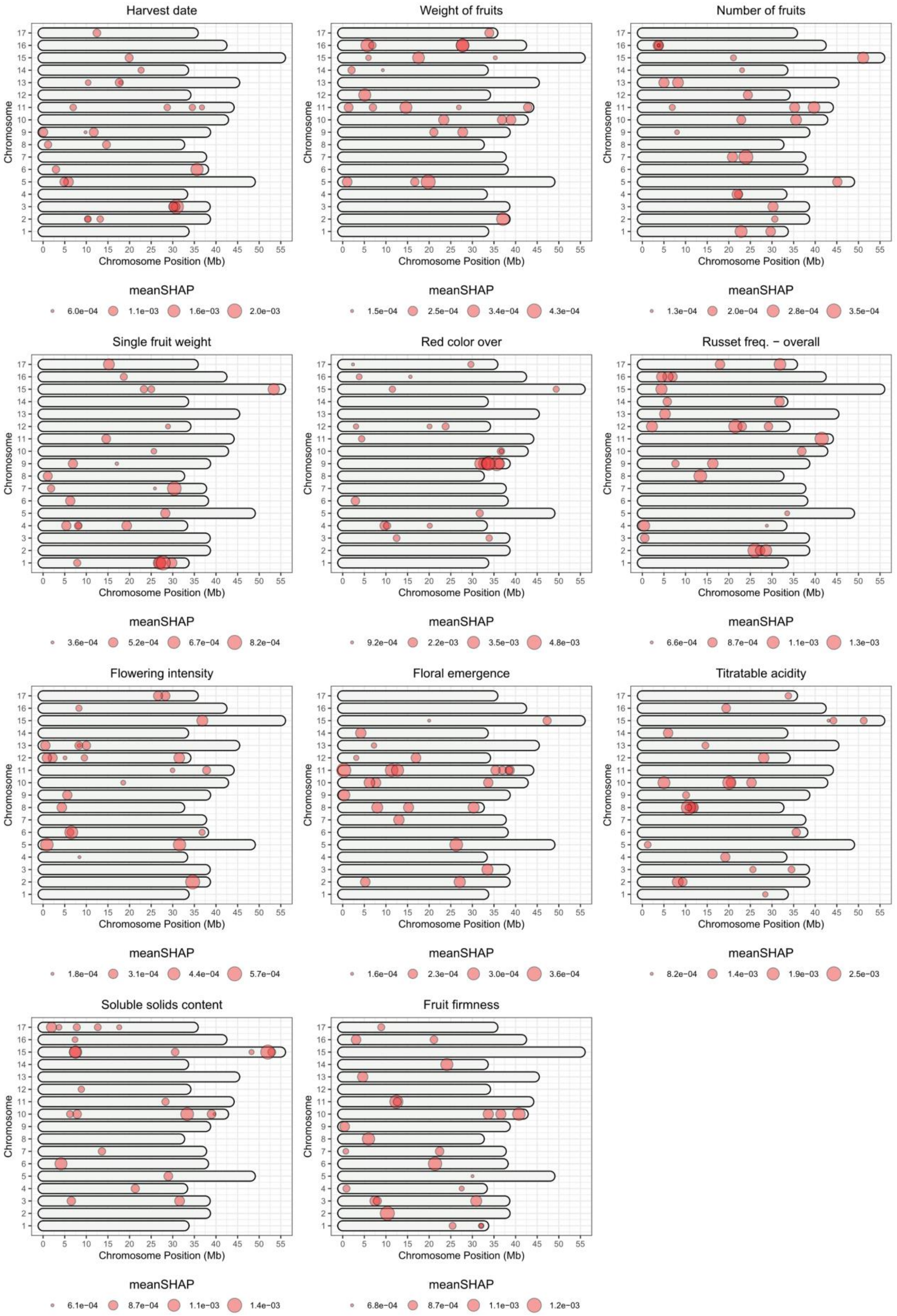


**Figure S5:** Chromosome positions for 25 SNPs with the highest absolute mean SHAP (Shapley additive explanations) found for each of the eleven studied traits.

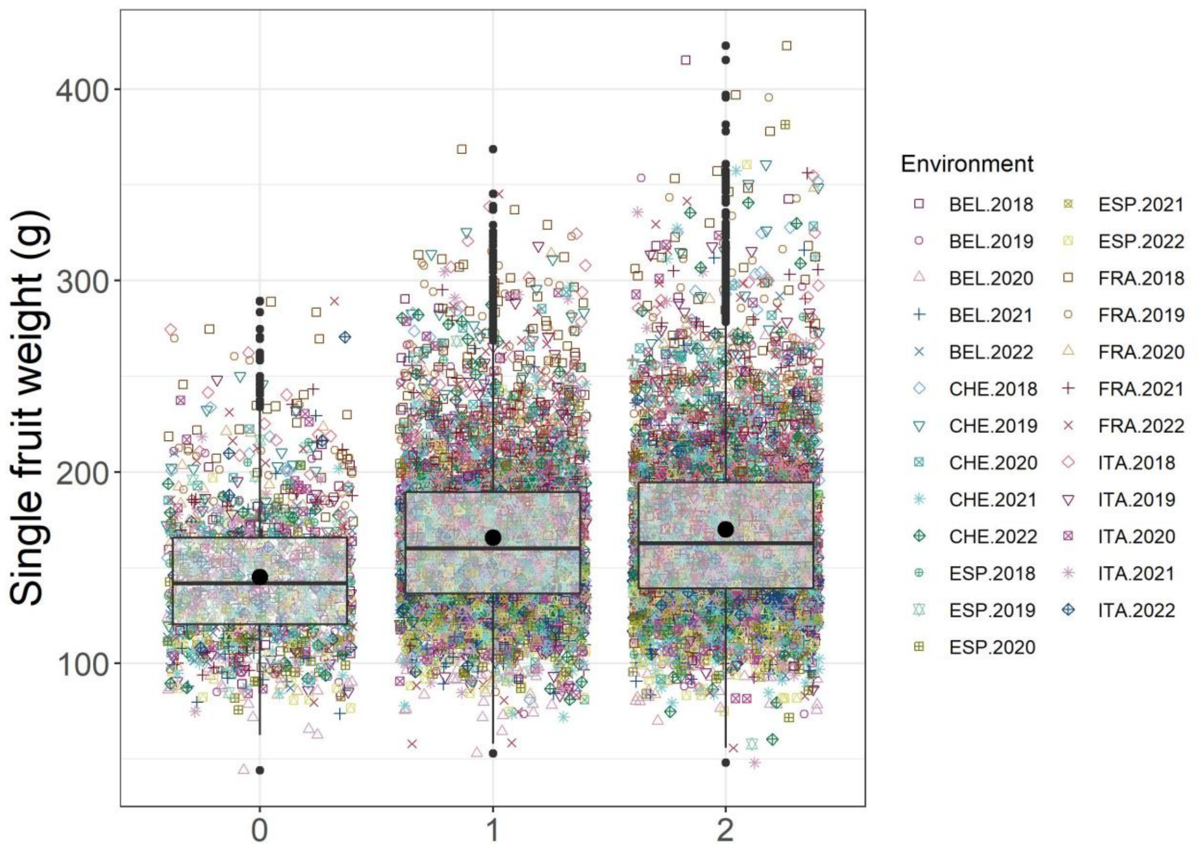


**Figure S6:** Comparison of adjusted means for single fruit weight across different allele dosages (0, 1, 2) of the marker AX-115591267 located on chromosome 1 at 28.6 Mb found by the deep learning approach. Black points indicate average values of adjusted means across environments: 145 g for allele dosage 0, 166 g for allele dosage 1, and 170 g for allele dosage 2.
